# Global organelle profiling reveals subcellular localization and remodeling at proteome scale

**DOI:** 10.1101/2023.12.18.572249

**Authors:** Marco Y. Hein, Duo Peng, Verina Todorova, Frank McCarthy, Kibeom Kim, Chad Liu, Laura Savy, Camille Januel, Rodrigo Baltazar-Nunez, Sophie Bax, Shivanshi Vaid, Madhuri Vangipuram, Ivan E. Ivanov, Janie R. Byrum, Soorya Pradeep, Carlos G. Gonzalez, Yttria Aniseia, Eileen Wang, Joseph S. Creery, Aidan H. McMorrow, James Burgess, Sara Sunshine, Serena Yeung-Levy, Brian C. DeFelice, Shalin B. Mehta, Daniel N. Itzhak, Joshua E. Elias, Manuel D. Leonetti

**Author notes:** co-first author (Marco Y. Hein, Duo Peng, Verina Todorova, Frank McCarthy, Kibeom Kim, Chad Liu). Max Perutz Labs, Vienna Biocenter Campus (VBC), Vienna, Austria.

## Abstract

Defining the subcellular distribution of all human proteins and its remodeling across cellular states remains a central goal in cell biology. Here, we present a high-resolution strategy to map subcellular organization using organelle immuno-capture coupled to mass spectrometry. We apply this proteomics workflow to a cell-wide collection of membranous and membrane-less compartments. A graph-based representation of our data reveals the subcellular localization of over 7,600 proteins, defines spatial protein networks, and uncovers interconnections between cellular compartments. We demonstrate that our approach can be deployed to comprehensively profile proteome remodeling during cellular perturbation. By characterizing the cellular landscape following hCoV-OC43 viral infection, we discover that many proteins are regulated by changes in their spatial distribution rather than by changes in their total abundance. Our results establish that proteome-wide analysis of subcellular remodeling provides essential insights for the elucidation of cellular responses. Our dataset can be explored at organelles.czbiohub.org.

## INTRODUCTION

Internal compartmentalization is an essential feature of human cellular organization. Within the cell, membrane-bound^1,2^ and membrane-less organelles^3,4^ provide spatial scaffolds that partition cellular functions, from oxidative ATP production in the mitochondrion to ribosome biogenesis in the nucleolus. This partitioning subdivides the cell into unique chemical environments that concentrate reactants and create the appropriate conditions for a wide range of biochemical activities^5^. Compartmentalization further enables protection from reactive intermediates and by-products (for example, peroxisomes segregate metabolic reactions that produce toxic oxidative species^6^) and provides a core mechanism to control cellular processes through the dynamic re-localization of cellular components^5,7^. The acquisition of intracellular organelles is the defining event in the evolution of eukaryotes^8^, equipping cells with drastically expanded capacities to remodel their composition and to interact with their environment.

Because of the central importance of spatial partitioning for cellular functions, defining the molecular composition of organelles and how this composition is remodeled across cellular states remains a major goal in human cell biology. As a result, multiple methods have been developed to define the proteomes of individual cellular compartments^9–11^. Intracellular localization can be sensitively and precisely characterized by microscopy imaging using immunofluorescence^12^ or the expression of fluorescent protein fusions^13,14^. However, these analyses are typically carried out serially using specific reagents or cell lines for each protein. As a result, proteome-wide coverage remains difficult to achieve with image-based approaches, which limits their applicability to analyze global cellular architecture and its remodeling. On the other hand, many strategies have been developed to characterize organellar proteomes and their dynamics using mass spectrometry^9,10^. Local protein neighborhoods can be defined by proximity ligation using organelle-specific markers tagged with labeling enzymes such as APEX^15^ or BioID^16–18^. The protein composition of entire cellular compartments can be further measured following immunoprecipitation^19–24^, biochemical fractionation or sequential solubilization methods^25,26^. Fractionation can be used to purify a specific organelle for in-depth analysis^27^, or to characterize all cellular compartments from a given sample using protein correlation profiling^28–35^. In correlation profiling, proteins from a given compartment distribute across several fractions in a manner that is distinct to that compartment^36^. Statistical methods are then used to cluster proteins that present similar enrichment patterns, thereby identifying proteins belonging to the same compartment.

Here, we present a new experimental and analytical strategy for the global characterization of the subcellular proteome. We use CRISPR/Cas9-based endogenous labeling to generate a library of HEK293T cell lines expressing epitope-tagged markers covering all major subcellular compartments, including membrane-less organelles. Our approach capitalizes on methods originally developed for the rapid immunoprecipitation (IP) of mitochondria^19,20^ and lysosomes^21^ following gentle cell homogenization. We couple organelle IP with liquid chromatography-mass spectrometry (LC-MS) to quantitatively define the subcellular proteome, totaling over 8,000 proteins. Our extensive coverage of subcellular structures combined with a graph-based analytical framework allows us to map protein localization on a global cellular scale (including many uncharacterized proteins), and to define the inter-connectivity network between individual compartments. Finally, we demonstrate how our method can be deployed to comprehensively profile dynamic proteome remodeling upon infection with the hCoV-OC43 β-coronavirus. Our results show that many proteins are regulated by changes in their spatial distribution rather than by changes in their total abundance, establishing that proteome-wide analysis of subcellular remodeling offers essential insights for the elucidation of cellular responses. Overall, we provide a combination of protocols, open-source code (<u>github.com/czbiohub-</u> <u>sf/Organelle_IP_analyses_and_figures</u>) and a web-accessible data resource (<u>organelles.czbiohub.org</u>) for the systems-level characterization of the human proteome’s architecture across cellular states.

## RESULTS

### Immunoprecipitation under native conditions provides rich organellar proteomes

Multiple studies have established that whole-organelle immunoprecipitation (IP) enables the analysis of native organellar proteomes, metabolomes and lipidomes by LC-MS^19–24^. In a native workflow, cells are mechanically homogenized under gentle conditions, exposing organelles that maintain their structural integrity and internal composition^20^. So far, pioneering studies have each primarily focused on a single membrane-bound organelle including mitochondrion^19,20^, lysosome^21^, peroxisome^22^, early-endosome^23^ and Golgi complex^24^. We reasoned that this strategy could be globally applied to all the different compartments of a given cell type to obtain a comprehensive map of the subcellular human proteome (Figures 1A-D). For each compartment, we used CRISPR/Cas9 to endogenously tag a reference protein marker (“bait”) with a cassette including a long amino-acid linker, GFP and a 3xHA epitope in HEK293T cells (Figure 1A). All markers were tagged on a cytosol-facing terminus, except for the plasma membrane markers for which an extracellular terminus was used. We used a long linker (102 amino-acids, Suppl. Figure S1A) to maximize exposure of the 3xHA IP epitope for capture. The addition of a fluorescent reporter (monomeric EGFP) enables the tagged cell lines to be used for imaged-based assays, while also providing a signal for the selection of engineered cells by flow cytometry. Markers for different organelles were tagged separately to generate an arrayed library of engineered cell lines (Figure 1B).

**Figure 1.**
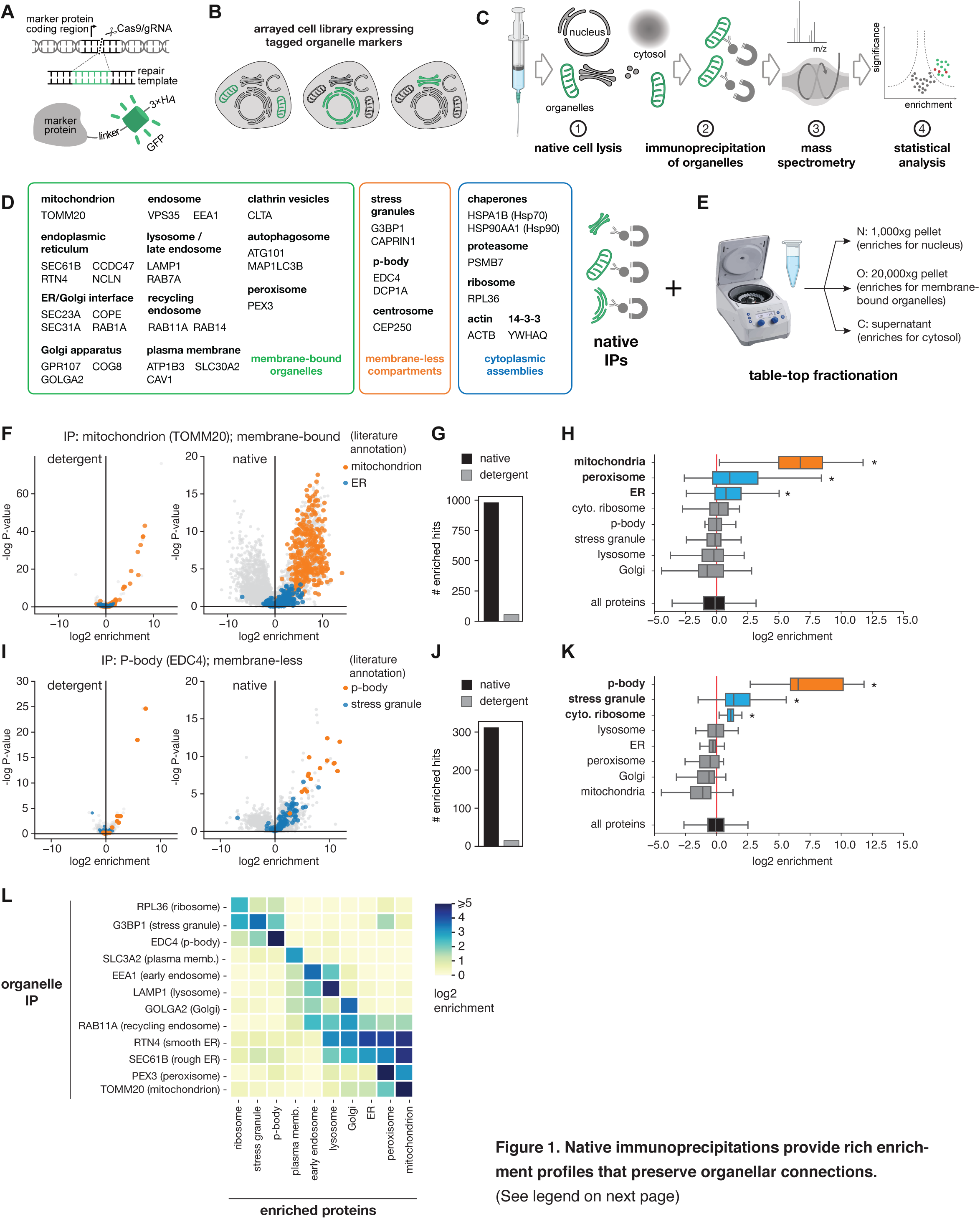
Native immunoprecipitations provide rich enrichment profiles that preserve organellar connections. **A)** CRISPR-based endogenous tagging of surface organellar markers in HEK293T cells. A 3xHA epitope is used for immunoprecipitation (IP). **B)** Markers for separate organelles are tagged separately to create an arrayed library of cell lines. **C)** Native immunoprecipitation workflow. Tagged cell lines are mechanically homogenized, exposing organelles that can be immuno-captured and analyzed by LC-MS. Protein abundance in each sample is then quantified for statistical analysis. **D)** List of 36 tagged proteins markers for a comprehensive coverage of subcellular compartments. **E)** Spin-fractionation provides enrichment for nuclear, organellar and cytosolic extracts. **F)** Protein enrichment profiles from detergent-based (1% digitonin) vs. native IP of TOMM20, a mitochondrial surface marker. Volcano plots show log2 enrichment of individual proteins in TOMM20 IPs relative to unrelated controls (See Material and Methods). P-values are calculated from a t-test using triplicate observations of each IP. **G)** Comparison of the number of significantly enriched hits (defined as log2 enrichment ζ1 and P-value ::0.01) for TOMM20 IPs in native vs. detergent-based conditions. **H)** Aggregate enrichment of proteins annotated to reside in different subcellular compartments in TOMM20 native IPs. In addition to mitochondrial proteins, peroxisomal and ER proteins are also significantly enriched (*: P ::0.05, t-test). **I-K)** Same as F-H, showing IPs for EDC4, a marker for p-body (membrane-less organelle). **L)** Heatmap showing log2 enrichments for proteins annotated to reside in different subcellular compartments (columns) across different native IPs (rows). Related to H and K.

For organelle isolation, native homogenates were prepared for each cell line by passing cells four times through the narrow opening of a 23G needle attached to a 1mL syringe, followed by rapid IP (10 min) of organelles with anti-HA magnetic beads and protein quantification by label-free LC-MS (Figure 1C, see Material and Methods). To obtain comprehensive coverage of different subcellular compartments, we applied this workflow to 36 separate protein markers localized to 14 membrane-bound or membrane-less organelles, as well as 5 cytoplasmic assemblies (Figure 1D). In addition, we used a simple three-step centrifugation protocol to enrich <u>N</u>uclear (1,000xg pellet), membrane-bound <u>O</u>rganellar (20,000xg pellet) and <u>C</u>ytoplasmic (supernatant) fractions from an untagged whole-cell homogenate (Figure 1E). These “N/O/C” fractions were included to capture nuclear and cytoplasmic proteins that would likely be excluded from the IP procedure, therefore helping us approach a comprehensive coverage of cellular composition. All IPs were performed in triplicate using 10^7^ cells per replicate harvested from 10cm-plate cultures.

Quantitative analysis revealed rich protein enrichment profiles for each triplicate organelle IP (Figures 1F-K; Suppl. Figure S2; see also Material and Methods). For example, IP of the mitochondrial outer-membrane translocase component TOMM20 showed enrichment for 982 proteins (enrichment ≥2-fold at p-value :s0.01 across triplicate experiments; Figures 1F-G), with the largest enrichment observed for proteins annotated as mitochondrial in the literature (Figure 1H; annotations from Suppl. Table 1). This was in stark contrast with the enrichment profile obtained for a TOMM20 IP obtained in mild detergent-containing buffer (Figure 1G, data from the OpenCell resource^14^). These results demonstrate the utility of detergent-free lysis for the native capture of cellular compartments. Native TOMM20 IP showed enrichments for proteins across all mitochondrial sub-structures (outer and inner membranes, inter-membrane space and matrix; Suppl. Figure S1C), verifying that our workflow preserves the structural integrity of organelles. Similar results were obtained for ER and lysosome IPs, showing comparable enrichment for membrane and lumenal components (Suppl. Figure S1D). In addition to preserving the integrity of individual organelles, our data also suggested that some contacts between organelles are maintained under native IP conditions. Indeed, TOMM20 IP also showed significant enrichment for peroxisomal or ER proteins, two organelles know to make direct contacts with the mitochondria^37,38^, but not for unrelated organelles such as lysosome or Golgi apparatus (Figure 1H). Strikingly, similar results were obtained for the IP of membrane-less compartments such as p-body and stress granules. For example, IP of the p-body marker EDC4 showed a strong enrichment for known p-body proteins, and weaker but statistically significant enrichment for proteins of stress granules or the cytosolic ribosome, two compartments known to directly interact with p-bodies^39^ (Figures 1I-H). Overall, mapping the global landscape of co-enrichment across all organelles revealed connections between interacting cellular compartments (Figure 1L).

### Graph-based analysis provides subcellular resolution to human proteome maps

We performed 58 triplicate IPs from our collection of 37 unique organelle markers (16 IP markers were profiled more than once). Our final dataset comprises a total of 8,195 unique proteins (“preys”) detected in at least one triplicate IP (see Material and Methods), and 8,541 total proteins isoforms. Each protein is defined by its enrichment across all 58 triplicate IPs, as well as by its normalized abundance in the N/O/C spin fractions (Figure 2A). This constitutes for each protein a 61-dimensional organellar enrichment profile that is closely related to its localization to specific cellular compartments. Indeed, enrichment profiles were highly correlated for proteins annotated in the literature to reside in the same organelle, and less correlated for proteins from separate organelles (Figure 2B, showing pairwise Pearson correlations; annotations from Suppl. Table 1). Reflecting the preservation of inter-organellar connections in the native IPs, enrichment profiles were also more highly correlated for proteins residing in compartments known to directly interact in the cell (for example between the ER and mitochondria, off-diagonal signal in Figure 2B). Furthermore, protein pairs engaged in high-stoichiometry interactions (for example, subunits of the same protein complex annotated in CORUM^40^) exhibited a strong correlation between their organellar enrichment profiles (Suppl. Figure S3A), mirroring the intracellular co-localization of strongly interacting proteins^14^.

**Figure 2.**
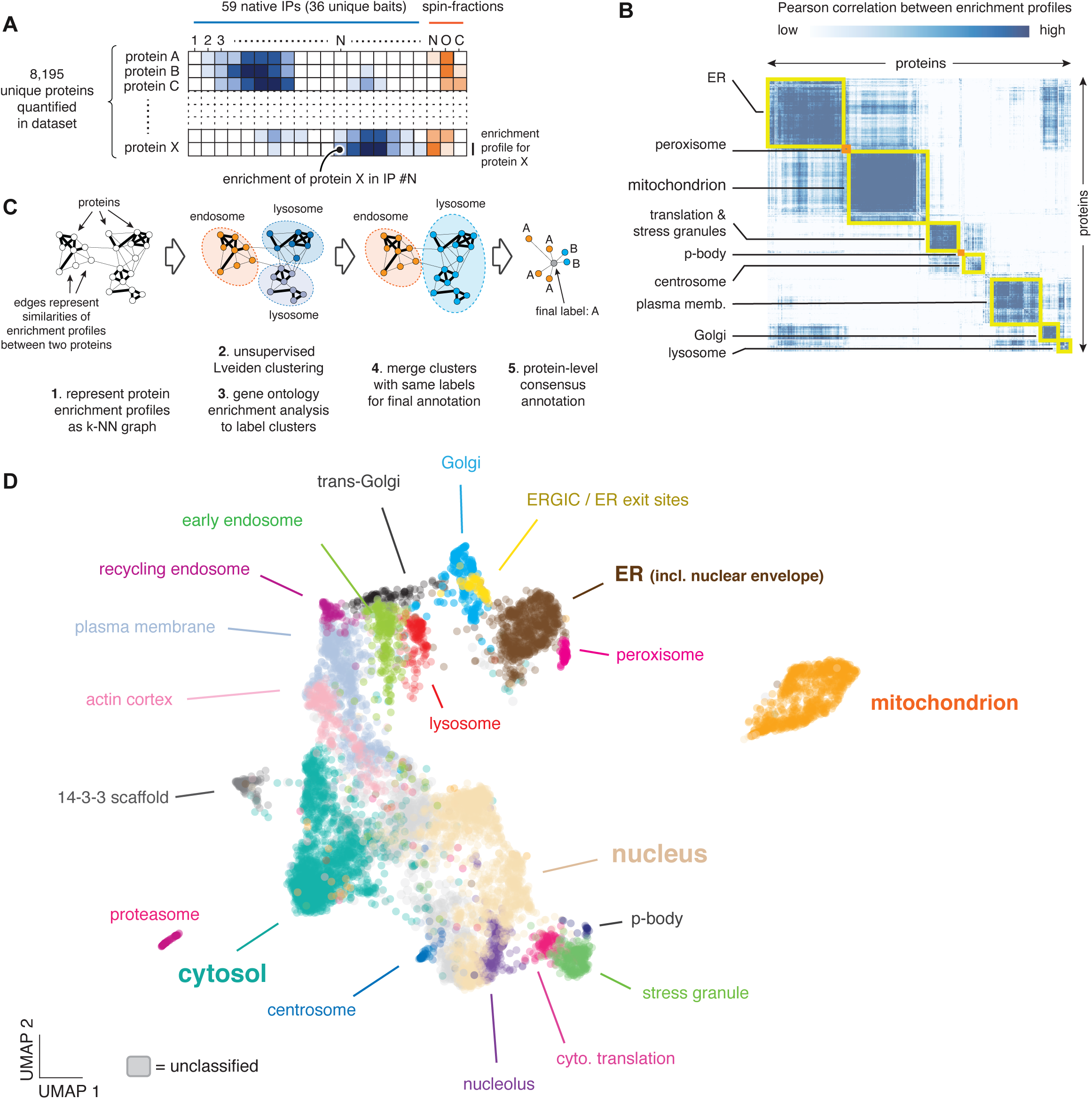
Graph-based analysis provides a subcellular map of the human proteome. **A)** Overall data structure. Our dataset identifies 8,195 unique proteins, each defined by its enrichment across individual native organelle IPs and spin fractions. The resulting enrichment profile vector provides a signature of the subcellular environment of each protein. **B)** Pairwise Pearson correlation matrix between enrichment profiles or annotated marker proteins. High correlation is observed between proteins annotated to reside in the same compartment (yellow highlights), and to a lesser extent between proteins residing in compartments known to interact in the cell (for example, ER and mitochondrion). **C)** Graph-based strategy for data-driven annotation of protein localization. The enrichment matrix is transformed into a k-nearest neighbor graph (1), from which clusters of co-enriched proteins are identified using Leiden clustering (2). Gene ontology enrichment analysis allows the annotation of clusters corresponding to separate subcellular compartments (3, 4). A final regularization step assigns to each protein the label most represented amongst its direct neighbors (consensus annotation; 5). **D)** Two-dimensional UMAP showing the 21 subcellular compartments identified by the enrichment analysis in C).

To further exploit the rich subcellular signatures contained in each protein’s enrichment profile, we developed an analytical strategy inspired by single-cell RNA sequencing (scRNASeq) workflows. In scRNASeq analysis, the relationship between individual cells is typically represented in a *k*-nearest neighbor (k-NN) graph in which edges correspond to the similarity of two individual cells in gene expression space, keeping edges with a number *k* of nearest neighbors for each cell. This graph representation is the foundation for dimensionality reduction algorithms such as UMAP^41^ and, separately, for unsupervised clustering using the Leiden algorithm frequently used for the annotation of cell types^42^. The ubiquity of k-NN-based analysis in scRNASeq reflects the power of graph-based representation to encapsulate complex relationships from high-dimensional data^43^. Therefore, we reasoned that a k-NN graph would be a powerful tool to represent the relationships between individual proteins’ enrichment profiles in our dataset.

We constructed a graph connecting each of the 8,541 proteins to its 20 nearest neighbors, with edges weights derived from the Euclidean distances between two proteins in the 61-dimensional enrichment space described above (Figure 2C; see Material and Methods). To identify proteins belonging to the same subcellular compartment, we performed unsupervised graph-based clustering using the Leiden algorithm^42^. We then objectively labeled each protein cluster according to its most enriched gene ontology term using the COMPARTMENTS^44^ and GO-*Cellular component*^45^ reference annotation databases, and merged clusters sharing the same ontology label. We finally refined these cluster-level annotations using a local neighborhood consensus strategy for individual proteins: a protein is annotated with the cluster-based label most found amongst its direct neighbors (Figure 2C). This consensus-calling step acts to better define boundaries between clusters. The complete enrichment and annotation data can be found in Suppl. Table 2. No definitive ontology-based annotation could be obtained for a total of 875 proteins (10% of the dataset). These mostly correspond to soluble proteins that lack a clear enrichment in N/O/C spin fractions (Suppl. Figure S3B), reflecting a broad multi-localization between nuclear, cytosolic and membrane compartments.

The result of our graph-based annotation analysis is visualized in Figure 2D as a two-dimensional UMAP, representing a spatial map of the HEK293T proteome (see also Suppl. Table 3 and <u>organelles.czbiohub.org/Subcellular_umap</u>). Overall, our graph-based method annotated the subcellular localization of 7,666 proteins. We distinguished 20 different protein clusters mapping to separate subcellular compartments, with sizes ranging from 18 (p-body) to 1,836 proteins (cytosol) (Suppl. Figure S4A). The arrangement of compartments within the map follows known biological relationships: for example, membrane-bound organelles are arranged in a sequence that matches the cell’s secretory pathway from ER to Golgi to endosomes to plasma membrane (Figure 2D). This suggests that our data can be analyzed to study the connectivity between compartments, which we explore in more detail below. In addition to classically defined compartments, our annotations also include a well-defined cluster of 130 cytosolic proteins defined by their strong enrichment in IPs of the 14-3-3 protein YWHAQ, which we labeled “14-3-3 scaffold”. 73% of proteins in that cluster were also identified in a recent proteomics study of 14-3-3 clients in HEK293T cells, supporting the annotation^46^.

To validate our annotations using an orthogonal approach, we trained a supervised machine learning classifier to predict the organelle membership of individual proteins based on literature-curated reference data (Suppl. Table 1). We used XGBoost^47^, a classifier based on gradient boosting^48^ that combines multiple decision trees to improve prediction performance. We limited classification to 15 compartments for which a critical mass of reference data was directly available (excluding, for example, trans-Golgi or recycling endosome, which most other datasets do not distinguish). For these 15 compartments, we performed receiver operating characteristic (ROC) analyses to quantify the agreement between graph- and classifier-based annotations (Suppl. Figure S4B). High values of area under the ROC curves (>0.94) were obtained in all cases, supporting our specific graph-based annotations and our overall computational approach.

Finally, we compared our annotation performance with that of other large-scale mapping studies. Such analysis is inherently difficult because no data exists that defines an objective ground truth; indeed, the agreement between different large-scale studies tends to be poor ^12,25,34,35,49^. The mitochondrion stands out, however, as a compartment for which reliable ground truth data is available. Given mitochondria’s central importance to cellular physiology, extensive efforts have been developed to precisely curate mitochondrial proteins. In particular, the MitoCarta^50^ (v3.0) and MitoCoP^27^ datasets combine large multi-modal data sources across cell types and cell states to define the mitochondrial proteome. We reasoned that these datasets could be combined to define a reference ground truth against which the annotation accuracy of large-scale mapping efforts could be objectively evaluated. Using this ground truth, we performed a precision/recall analysis on the lists of mitochondrial proteins annotated in recent proteome-scale datasets from a diversity of mass spectrometry or imaging approaches (Suppl. Figure S4C). Our annotations achieved the highest recall across all datasets (77%, 1.45x higher than the average recall), while being one of only two datasets to exhibit a false-discovery rate lower than 5% (3.1%). This analysis of mitochondrial annotations, together with the classifier results presented above, establish that our experimental and analytical strategy offers a robust method for the definition of the subcellular proteome.

### Subcellular protein networks define functional signatures and quantify cellular organelle connections

The k-NN graph representation described above (subsequently referred to as the “k-NN map”) contains information that extends beyond organellar membership. Indeed, a given protein’s local network in the k-NN map provides a specific signature that can help define its cellular function. Figures 3A and 3B show the example of the local k-NN network centered on WASHC5 (Strumpellin), a core component of the actin-nucleating WASH complex (WASHC). WASHC is a key regulator of endosomal trafficking that modulates the transport of protein cargo from endosomes to other compartments including trans-Golgi network (TGN), plasma membrane and lysosomes^51–53^. In the k-NN map, WASHC5 localizes at the nexus between endosomal, TGN and lysosomal clusters (Figure 3A), reflecting its cellular function. More specifically, WASHC performs its role by associating separately with two other complexes: Retromer and Commander (retriever/CCC)^53–55^. Retromer and Commander both control endosomal tubulation but regulate the trafficking of different sets of cargoes. Recapitulating these complex functional relationships, WASHC subunits are connected to both Retromer and Commander in the k-NN map, but Retromer and Commander occupy distinct territories (Figure 3B). Another example is shown in Suppl. Figure S5, presenting the local k-NN network centered on NUP205, a component of the nuclear pore complex. NUP205 is positioned near the interface between nuclear and ER clusters, mirroring the contiguity of the nuclear envelope with the ER. Furthermore, the NUP205 k-NN neighborhood includes many nuclear pore subunits, as well as key components of the nuclear lamina and of the nuclear import machinery (see annotations in Suppl. Figure S5A). Altogether, the WASHC5 and NUP205 examples illustrate how the detailed structure of the k-NN map can be mined for functional exploration. To facilitate such analysis, all k-NN connections between individual proteins are compiled in Suppl. Table 4 and can also be explored online at <u>organelles.czbiohub.org/Protein_network_graph</u>.

**Figure 3.**
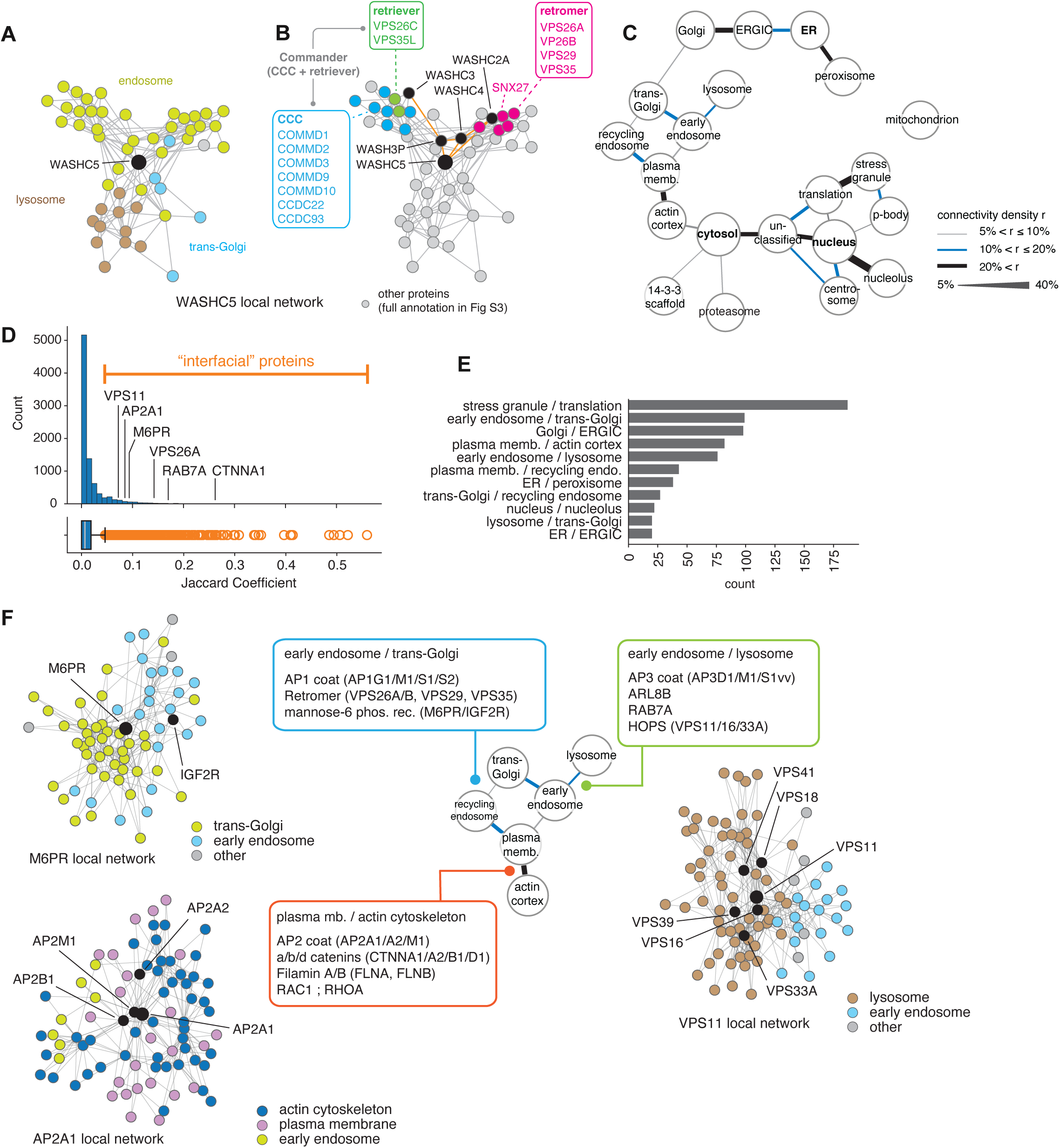
Protein spatial networks identify co-functioning proteins and organelle interfaces. **A)** Local k-NN network centered on WASHC5, a subunit of the WASH complex that regulates protein trafficking between endosomes, lysosome and trans-Golgi network. **B)** Annotated WASHC5 k-NN network, highlighting individual subunits for the WASH, Retromer and Commander (retriever and CCC) protein complexes. **C)** Density of k-NN connectivity between organelle clusters quantifies inter-compartment crosstalk. In this visualization, annotation clusters are arranged to replicate the arrangement in Figure 2D. **D)** Interfacial proteins are defined as high-value outliers in the distribution of Jaccard coefficient in the entire dataset. Bottom: boxplot showing the 25th, 50th, and 75th percentiles of the distribution; whiskers represent 1.5 times the interquartile range. **E)** Main organelle/organelle interfaces represented in the set of interfacial proteins. **F)** Examples of interfacial proteins in the vesicular pathway. See text for details.

The k-NN map can also be exploited to quantify the connectivity between organelles. We measured the inter-organelle connectivity density ρ defined by the number of k-NN connections bridging proteins from two separate annotation clusters, normalized by the size of each cluster. A graph visualization of the strongest connectivity densities (ρ > 5%) revealed a network reflecting known functional relationships between organelles, with the strongest connections bridging ERGIC/Golgi, plasma membrane and actin cortex, or the translation machinery and stress granule (Figure 3C).

Inter-organellar communication is an essential part of cellular physiology^2^; thus, many proteins are specialized to act at the interface between organelles, ensuring the transport of molecules, the propagation of cellular signals or the coordination of metabolic pathways. To further explore organellar connectivity in our dataset, we identified “interfacial” proteins in the k-NN map. We adopted a modified version of the Jaccard coefficient, a measure used in graph theory to quantify a given node’s propensity to make connections with two separate clusters (see Material and Methods). Therefore, we quantified the distribution of Jaccard coefficient for all proteins in the k-NN map using graph-based annotation clusters (Figure 3D-E). Values were low for most proteins (median = 0.0075), but the distribution showed a long tail of proteins with higher Jaccard coefficients (Figure 3D). We defined interfacial proteins as the high-value outliers in this distribution (values above third percentile + 1.5x inter-quartile range; Figure 3D). The corresponding list of 950 interfacial proteins (Suppl. Table 5) shows an over-representation for interfaces between organelles sharing high connectivity densities (e.g., Golgi/ERGIC or plasma membrane/actin cortex, compare Figures 3C and 3E). This list includes 187 proteins at the interface between the translation machinery and stress granules, reflecting the key role that stress granules proteins play in the regulation of translation^56^.

Figure 3F presents a more detailed exploration of interfacial proteins within the vesicular system. Our data-driven analysis correctly identifies many protein systems known to specialize in transport or regulation at specific organellar boundaries. These include AP1, AP2 and AP3 coat proteins, which scaffold clathrin-mediated transport at the endosome/TGN, plasma membrane/endosome/actin cortex and endosome/lysosome interfaces, respectively^57,58^ (Figure 3F, all panels). At the plasma membrane, the actin cortex plays a central role for trafficking endocytic plasma membrane cargo towards the endosomal system^58^, which is mirrored by the position of the AP2 coat complex in the k-NN map (Figure 3F, bottom left). Other examples include: 1) the M6PR and IGF2R mannose-6-phosphate receptors^59^, and the retromer complex^53^ which mediate recycling between endosome and trans-Golgi (Figure 3F, top left); 2) at the endosome/lysosome interface, the HOPS complex^60^ which tethers endosomes and lysosomes, and the small GTPases ARL8B^61^ and RAB7A^62^ (Figure 3F, right); 3) the catenin^63^ (CTNN) and filamin^64^ (FLN) protein families that play a key role in bridging the actin cytoskeleton to plasma membrane effectors, an interface that also includes the small GTPases RAC1 and RHOA which regulate membrane actin polymerization to control cell migration^65^ (Figure 3F, bottom left). Overall, these results demonstrate how our dataset can be leveraged for the analysis of inter-organellar communication.

### Annotating protein subcellular localization

Our protein-level annotations of subcellular localization are summarized in Supplementary Table 6. We report results from the graph-based clustering analysis described above (including interfacial proteins), together with high-confidence predictions from the XGBoost machine learning classifier (reporting predictions for which classification probability exceeded 80%). The two annotation strategies provide complementary insights for the comprehensive curation of subcellular protein localization. Graph-based methods that rely on unsupervised clustering enable *de novo* annotations, even when no critical mass of reference data is available to train a classifier. On the other hand, classifier-based methods might be more precise in cases where sufficient ground truth is available. Using the mitochondrial proteome as a point of reference, we observed that graph-based annotations supported a higher recall than classifier-based annotations (77% vs 70%), at the expense of a slight increase in false-discovery rate (3.1% vs 1.2%). A defining advantage of graph-based methods is to provide an explicit framework in which to identify proteins acting at the interface between cellular compartments. This produces an additional layer of annotation that cannot be captured by curation that only assigns a single label for each protein. Our final annotation set includes graph-based annotations for 7,666 proteins (90% of the dataset) and high-confidence classifier-based annotation for 6,502 proteins (76% of the dataset). Across the 15 compartments defined in both sets, graph- and classifier-based annotations shared 90% matching annotations (Suppl. Figures S4D-E).

### De-orphaning protein localization

Our annotations include many proteins for which no localization information exists in reference databases. To further validate these annotations, we performed a microscopy-based analysis to de-orphan the localization of poorly characterized membrane proteins. We first curated an unbiased list of transmembrane or lipidated proteins for which no subcellular localization was annotated in reference databases (Figure 4A). Our dataset provided localization annotations for 1,237 membrane proteins in HEK293T cells. We identified “orphan” proteins from this list by filtering for proteins with minimal annotation in the COMPARTMENTS dataset^44^ (256 proteins of score ::3, indicating low-confidence or absent annotations) that also had no “Subcellular location” specified in Uniprot^66^. This established a final set of 82 membrane proteins of unknown or poorly characterized subcellular localization. We then used CRISPR/Cas9 to endogenously tag all 82 proteins with a split-mNeonGreen fluorescent cassette separately at their N- or C-terminus. 31 tagged cell lines exhibited enough fluorescent signal to be selected by flow cytometry and serve as a final set for our analysis. We characterized the subcellular localization of all 31 orphan proteins using confocal fluorescence microscopy in live cells (Figure 4B). In 29 out of 31 cases, the subcellular distribution of the tagged protein matched our annotations (90%, Figure 4C). These results further validate our experimental and analytical strategy for the prediction of protein localization in the human cell. Among the two cases for which imaging results did not match the protein target’s annotated localization, TPRA1 displayed a non-uniform distribution at the lysosomal surface while being annotated as interfacial between trans-Golgi and recycling endosome in our proteomics dataset. The other case, GLT8D1, presented a nucleolar localization by imaging, while being annotated as a Golgi/trans-Golgi protein in our proteomics dataset. A nucleolar localization would be unexpected for a type-II transmembrane protein such as GLT8D1. Indeed, a literature search uncovered that GLT8D1 has been previously found to localize to the Golgi and trans-Golgi in human cells^67,68^. Therefore, we hypothesize that the nucleolar localization we observe might be a tagging artifact.

**Figure 4.**
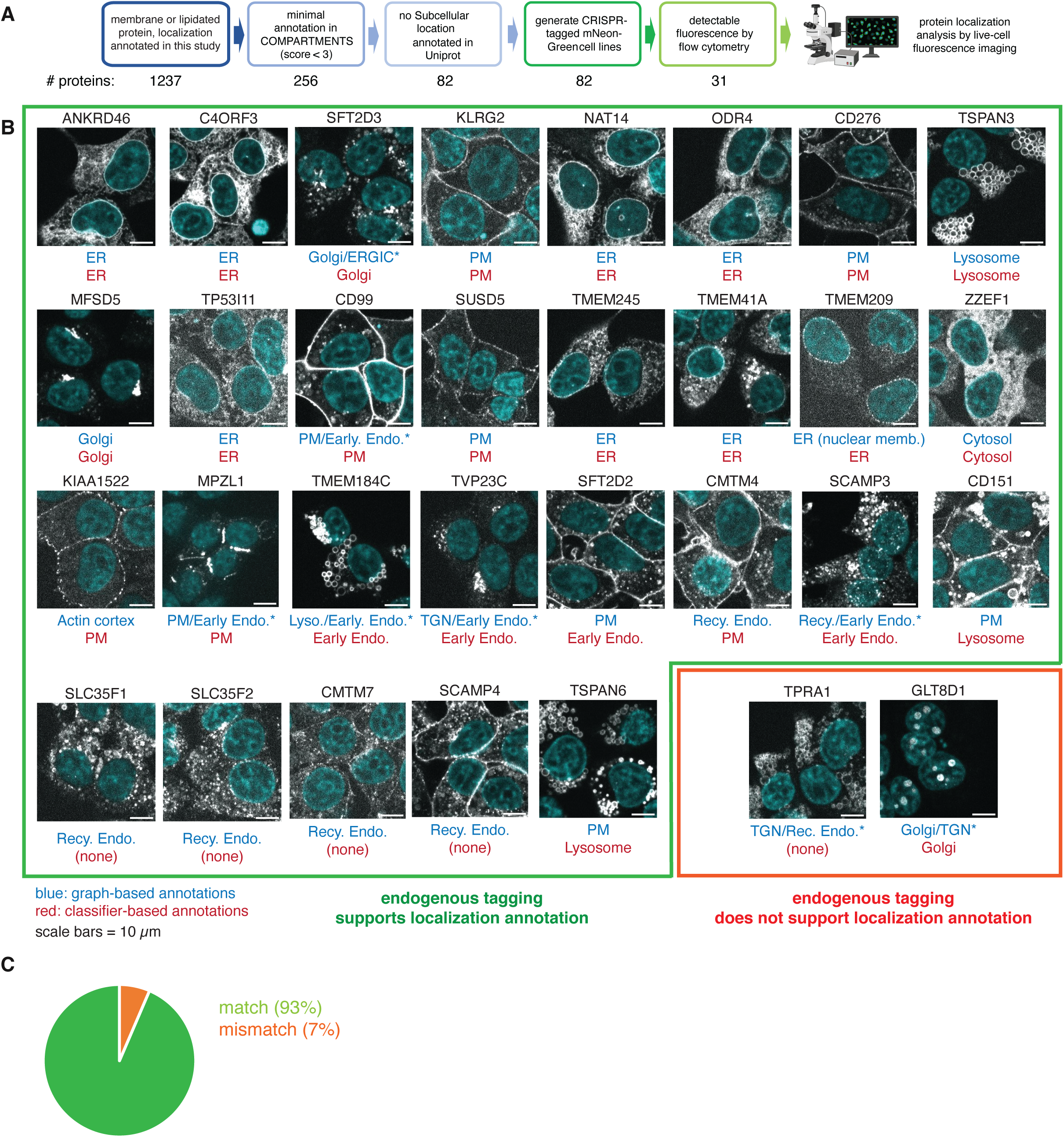
De-orphaning subcellular localization. **A)** Strategy to identify and characterize proteins with minimal annotations for subcellular localization in public databases. A final 31 protein targets were characterized. **B)** Representative images showing the subcellular localization of the 31 proteins targets, endogenously tagged with a split-mNeonGreen system in HEK293T and characterized by spinning disk confocal fluorescence microscopy. Graph-based (blue) and classifier-based (red) localization annotations are reported. Asterisks denote interfacial proteins. **C)** Aggregate results: endogenous tagging supports the annotations of subcellular localization in 93% of cases (29 out of 31 targets).

### Pan-cellular remodeling during hCoV-OC43 infection

Altogether, our organelle profiling strategy provides a framework to extract global features of intracellular organization. We next applied this strategy to capture the intracellular remodeling at play during cellular perturbation by profiling cells infected with the coronavirus hCoV-OC43, a common respiratory pathogen typically causing mild cold-like symptoms in humans^69^. OC43 is an enveloped, positive-sense single-stranded RNA virus that belongs to the same β-coronavirus family as SARS and SARS-CoV-2 but only requires biosafety level-2 containment. We infected HEK293T at a multiplicity of infection (MOI) of 0.25 and profiled their proteome 48 hours post infection (48 hpi). Over 80% of cells were infected at this time point as measured by immunofluorescence (Suppl. Figure S6A). We selected a subset of 25 tagged cell lines (Suppl. Figure S6B) and performed organelle immunoprecipitation and N/O/C spin fractionation in uninfected control vs. infected conditions, followed by proteomic quantification by LC-MS. In each condition, the enrichment matrix for all proteins was translated into a k-NN graph for analysis (Figure 5A). To transform k-NN graphs into a representation from which distances can be calculated, we used the aligned-UMAP algorithm^41^ to project graph data into a shared 10-dimensional Euclidean space (See Material and Methods). We finally profiled the movement of individual proteins in this aligned space by measuring the Euclidean distance between their infected vs. uninfected vectors (Figure 5A, right). For each protein, this distance represents a subcellular remodeling score that measures that protein’s repositioning in the underlying cellular k-NN map. The distribution of remodeling scores across the entire dataset showed that most proteins had low scores (indicating an overall conservation of their k-NN neighborhoods), with a long tail of high-value outliers (Figure 5B; outliers defined above third percentile + 1.5x inter-quartile range). These 633 outliers define a group of “infection hits” (Suppl. Table 7): proteins whose subcellular environment changes significantly during infection. The 633 infection hits encompass 8% of the proteome measured in our dataset, indicating the large extent of pan-cellular remodeling occurring upon viral infection. To visualize how individual infection hits distribute across subcellular compartments, we produced aligned two-dimensional UMAPs under uninfected and infected conditions. To facilitate visualization, we performed low-resolution Leiden clustering to separate the UMAPs into 10 broad subcellular territories (Figures 5C-E; Suppl. Table 7). Infection hits were found in all territories and exhibited a wide variety of remodeling behaviors (Figure 5E, diagonal trajectories). A movie of morphed trajectories between the two conditions captures the remodeling behavior of individual proteins (Suppl. Movie 1).

**Figure 5.**
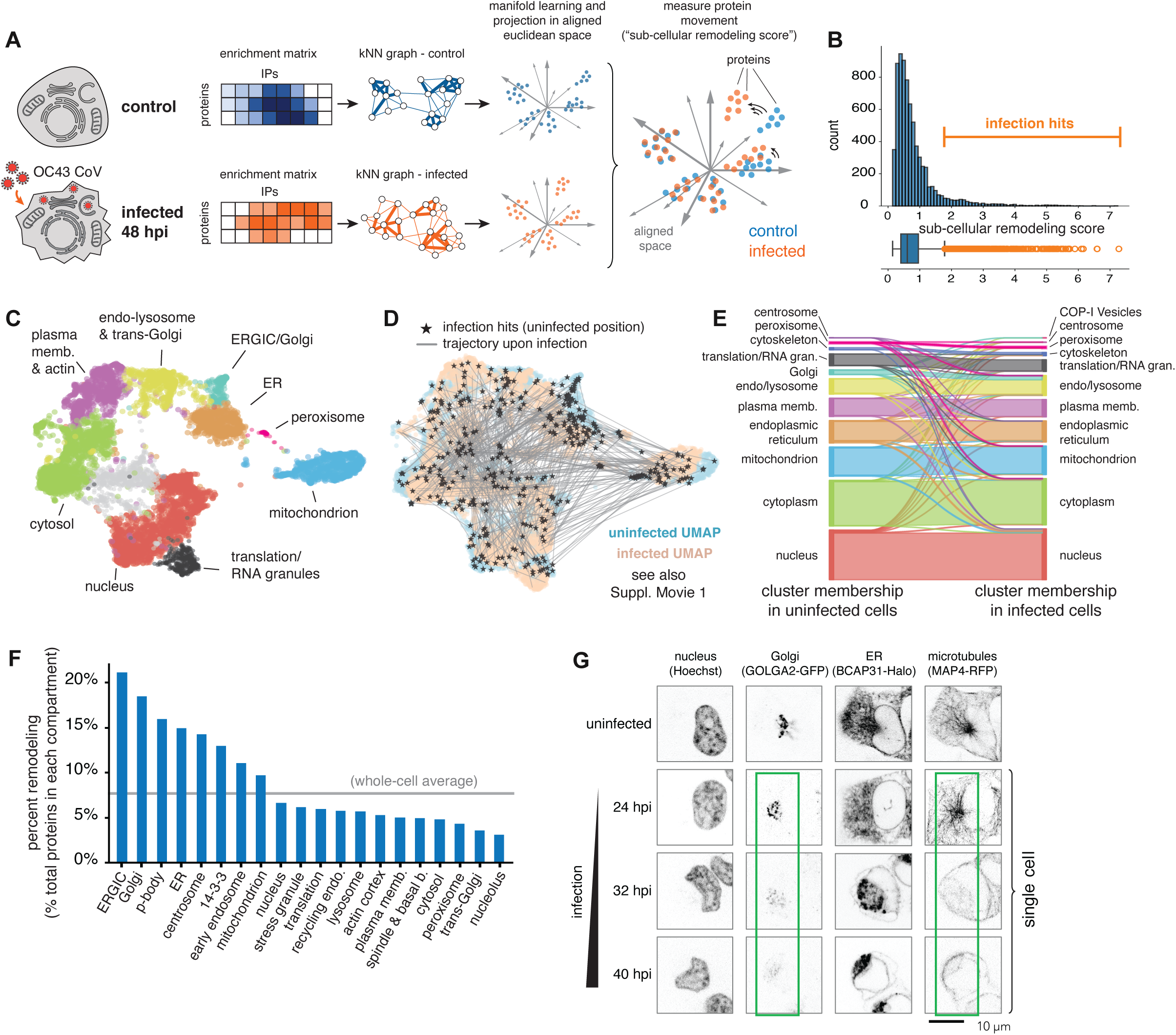
Pan-cellular remodeling during OC43 CoV infection. **A)** Data analysis strategy for the graph-based quantification of subcellular remodeling. A subcellular remodeling score is quantified for each protein, representing remodeling of local subcellular environment. See text for details. **B)** Distribution of subcellular remodeling scores across the dataset. Interfacial proteins are defined as high-value outliers in the distribution. Bottom: boxplot showing the 25th, 50th, and 75th percentiles of the distribution; whiskers represent 1.5 times the interquartile range. **C)** Two-dimensional UMAP of uninfected organelle profiling dataset showing subcellular territories defined by low-resolution Leiden clustering. **D)** Aligned uninfected (blue) and infected (salmon) 2D-UMAPs showing individual 2D trajectories for infection hits. The original position of each hits in uninfected conditions is marked by a black star. Related to C). **E)** Sankey diagram showing the distribution of infection hits (diagonal lines) across subcellular territories in uninfected (left) and infected (right) conditions. Related to C) and D). **F)** Proteome remodeling within individually resolved compartments, showing the fraction of infection hits over total number of proteins in each compartment. **G)** Live-cell confocal microscopy imaging from endogenously tagged cell lines shows significant remodeling of Golgi, ER and microtubule network.

Quantifying the fraction of infection hits within individual organelles provides a finer picture of subcellular re-organization (Figure 5F, using graph-based reference annotations defined above). ERGIC, Golgi, centrosome and ER exhibited some of the highest degrees of remodeling (Figure 5F). This matches the known cell biology of coronavirus replication and assembly at the ER/Golgi interface: structural proteins are inserted into the ER membrane and transit to the ERGIC to reach complex replication/assembly organelles where mature virions are formed^70,71^. Viral particles then traffic to the Golgi apparatus for glycosylation and other post-translational modifications^71^. We verified the pronounced remodeling of the Golgi, ER and centrosome using live-cell microscopy. HEK293T cells were endogenously tagged with fluorescent reporters on three separate organelle markers: GOLGA2 (GM130, cis-Golgi; GFP tag), BCAP31 (ER; Halo tag) and MAP4 (microtubule cytoskeleton; RFP tag). Upon infection, we observed a striking fragmentation of the Golgi complex, accompanied by a condensation of the ER (Figure 5F). This was accompanied by large changes in the distribution of microtubules in infected cells, compatible with a loss of the microtubule organizing center following centrosomal remodeling. In SARS-CoV-2, the NSP13 viral protein has been shown to directly interact with host proteins involved in both centrosomal and Golgi organization, suggesting a possible link between the remodeling of both compartments^72^. While an in-depth elucidation of the mechanisms driving ER/Golgi remodeling upon OC43 infection is beyond the scope of the current study, we noticed that the distance between Golgi and lysosomal proteins in organellar enrichment space decreased significantly during infection (Suppl Figure S6C). This might mirror the unconventional lysosomal exocytosis route that has been proposed for the egress of β-coronaviruses, which is believed to involve direct protein transport from Golgi to lysosomes^71^.

### Subcellular remodeling reveals cellular responses not captured by abundance changes

In addition to organelle profiling, we measured whole-cell abundance changes of transcripts and proteins in OC43-infected vs. control conditions (Suppl. Figure S7A-B). We observed a lack of overlap between infection hits defined from subcellular remodeling and differentially abundant proteins or transcripts (Figures 6A-B; Suppl. Figure S7A-B). This suggests orthogonal modes of functional regulation: many proteins are dynamically regulated at the level of their spatial environment rather than by changes in cellular abundance. Vice versa, many proteins are regulated at the abundance level, while their subcellular niche stays the same. Analyzing the different groups of infection-regulated proteins or transcripts with pathway enrichment^73,74^ demonstrated that profiling subcellular remodeling enables the identification of cellular responses not captured by abundance changes (Figures 6A-B). Specifically, subcellular infection hits were enriched for proteins involved in the metabolism of glycosaminanoglycans (GAGs, a group of surface-exposed polysaccharides that includes heparan sulfate glycans), as well as in ferroptosis, autophagy and cellular senescence (Figure 6A). Multiple lines of evidence support the direct relevance of these responses to the cell biology of OC43 infection: 1) GAGs are the surface receptor for OC43 entry under cell culture conditions^75^, so that modulation of their metabolism (which remains poorly understood^76^) might be beneficial for the viral life cycle, or for innate immune responses ; 2) autophagy plays an important role in the replication and pathogenesis of multiple coronaviruses^77,78^, including OC43^79^; 3) senescence is a primary host cell response to infection by many viruses, for example SARS-CoV-2^80^.

**Figure 6.**
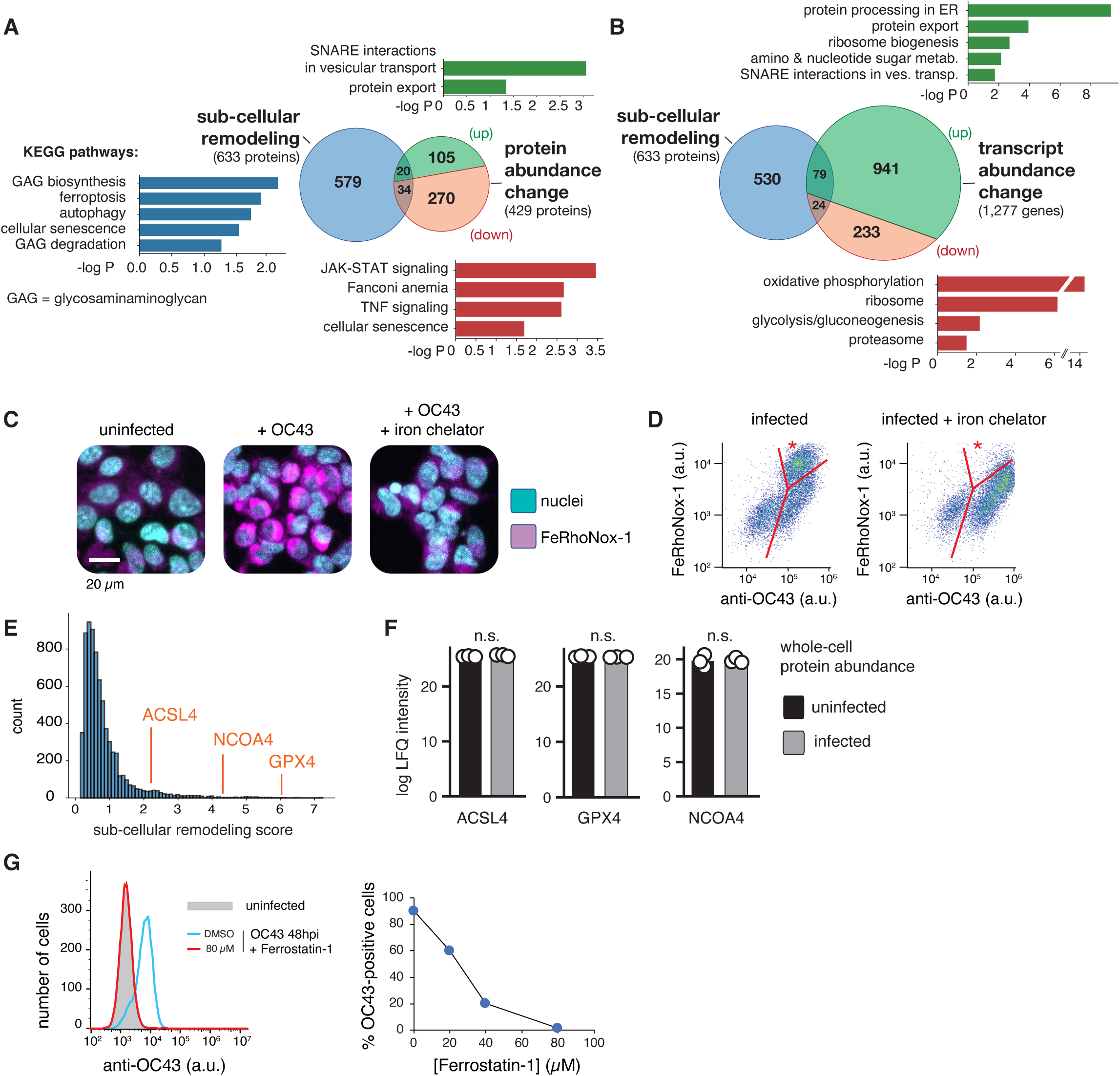
Subcellular remodeling upon OC43 infection reveals cellular responses not captured by abundance changes. **A)** KEGG pathway enrichment analysis of infection hits from subcellular remodeling (633 proteins, blue) vs. proteins that exhibit significant changes in whole-cell abundance (429 differentially abundant proteins total, including 125 up-regulated proteins, green, and 304 down-regulated proteins, red). P-values: t-test. **B)** Same as A), showing differentially regulated transcripts (1,277 differentially abundant proteins total, including 1,020 up-regulated transcripts, green, and 257 down-regulated transcripts, red). P-values: t-test. **C)** Ferropotosis induction in OC43-infected cells. Live cells were treated with 5 µM FeRhoNox-1 (magenta) to detect ferroptosis, and with 1 µg/mL Hoechst 33342 to stain nuclei (cyan). Nitroxoline (12 µM) was used to chelate Fe^2+^ ions. **D)** Flow cytometry quantification of OC43 protein levels (anti-OC43 monoclonal antibody 541-8F, exact epitope unknown) and FeRhoNox-1 (5 µM, ferroptosis reporter). Treatment with the iron chelator nitroxoline (12 µM) led to a significant decrease in the number of cells positive for both OC43 and FeRhoNox-1 (sub-population marked by an asterisk). **E)** Subcellular remodeling scores of ACSL4, NCOA4 and GPX4 (cf. Figure 5B). **F)** Whole-cell protein abundance measured by bulk proteomics. **G)** Detection of OC43-infected cells by flow cytometry (anti-OC43 monoclonal antibody 541-8F, exact epitope unknown). Treatment of cells with the ferroptosis inhibitor Ferrostatin-1 led to a decrease in OC43 infection.

We further explored ferropotosis because this response was recently implicated in the regulation of replication for OC43 and other positive-strand RNA viruses^81^. Ferroptosis is a form of non-apoptotic and non-necrotic programmed cell death dependent on cellular iron^82,83^. During ferroptosis, iron (primarily Fe^2+^) catalyzes unrestrained lipid peroxidation, leading to oxidative stress and to a gradual destruction of intracellular membranes^83^. Under normal conditions, intracellular iron is stored in ferritin, a high-molecular weight protein cage. The autophagy-mediated degradation of ferritin (ferritonophagy) leads to the release of free iron in the cytoplasm and acts as a major driver of ferroptosis^84–86^. Multiple independent protein pathways play a central role in the induction and regulation of ferroptosis in cells, including ACSL4 (an acyl-transferase that incorporates polyunsaturated fatty acids into lipids, generating the substrates for peroxidation^87^), GPX4 (a peroxidase that clears toxic peroxides, counteracting ferroptosis^83^) and NCOA4 (the autophagy adapter for ferritonophagy^88^).

We validated the induction of iron-dependent ferroptosis in our OC43 infection model: infection led to an increase of intracellular Fe^2+^ as detected by the fluorescent probe FeRhoNox-1, which could be reversed by the iron chelator nitroxoline (Figure 6C-D). Furthermore, ACSL4, GPX4 and NCOA4 all exhibited high subcellular remodeling scores (Figure 6E), suggesting that multiple ferroptosis pathways might be regulated upon OC43 infection. This remodeling was not accompanied by changes in protein abundance (Figure 6F). Strikingly, inhibition of ferroptosis with Ferrostatin-1 led to a significant decrease of OC43 infection in HEK293T (Figure 6G), as was recently demonstrated in monkey LLC-MK2 cells^81^. While follow-up work will be needed to better characterize the mechanistic interplay between ferroptosis and the viral life cycle, these results establish the key significance of ferroptosis in OC43 infection. Altogether, this analysis demonstrates how profiling subcellular remodeling can complement whole-cell proteomics or transcriptomics assays to capture the full landscape of cellular responses triggered by a given perturbation.

### Interactive data sharing at organelles.czbiohub.org

To facilitate access and exploration of our datasets, we built an interactive web application at <u>organelles.czbiohub.org</u>. This portal provides an intuitive interface to explore and download enrichment data from individual organelle IPs, protein-level k-NN networks, subcellular localization UMAPs, and subcellular remodeling induced by OC43 infection (summarized in Figure 7).

**Figure 7.**
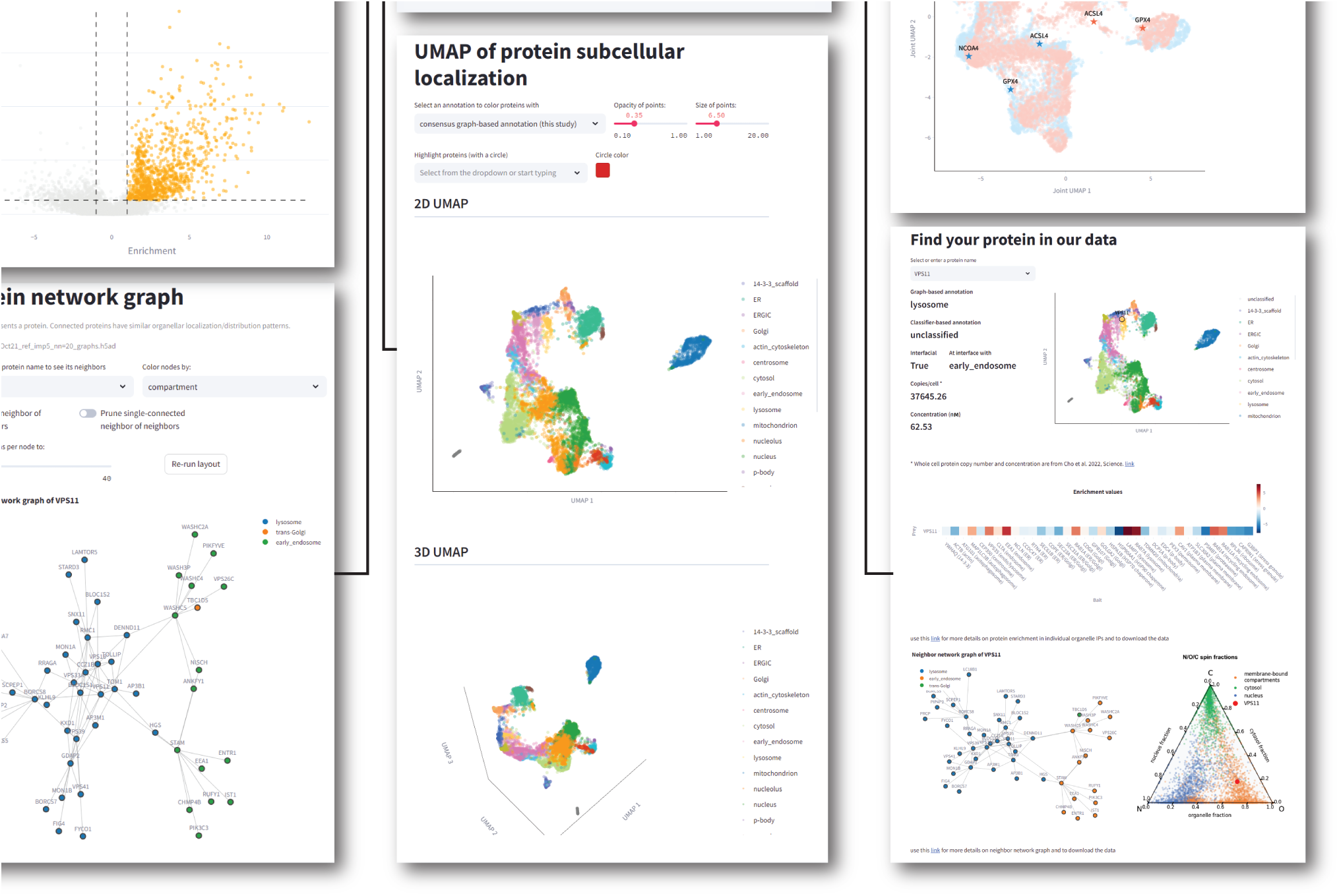
Interactive data exploration. Summary of the <u>organelles.czbiohub.org</u> web data portal. See website for details.

## DISCUSSION

Here, we described an integrated experimental and analytical framework to advance the characterization of the subcellular proteome and its dynamics. Our approach combines three main features. First, we applied native immunoprecipitation workflows to a cell-wide collection of membrane-bound and membrane-less markers. Second, we developed a graph-based formalism to define functional relationships between proteins, in particular to annotate their subcellular localization. Third, we extended this strategy to profile proteome dynamics and identify proteins whose subcellular environment gets remodeled during cellular perturbation. Applying our approach to characterize the cell biology of hCoV-OC43 infection, we demonstrated that profiling subcellular remodeling can reveal cellular responses not captured by abundance changes.

Our experimental workflow builds upon methods developed for the native isolation of organelles^19–24^. Expanding from previous studies, our strategy uses CRISPR/Cas9-based endogenous tagging, which circumvents the risk that over-expression might affect the enrichment of proteins in organelle pull-downs^20^. We also show that native isolation methods provide rich protein composition profiles for membrane-less organelles, demonstrating that this strategy can help characterize a wide range of cellular compartments. Finally, we applied native IP to an extensive collection of organelle markers to approach a comprehensive coverage of cellular architecture. As a result, our dataset provides for each protein a complex signature of enrichment across many different pull-downs. This extensive coverage is particularly beneficial in the context of native IPs that can preserve significant connections between organelles. A single IP might not provide enough information to clearly disambiguate ER vs. mitochondrial proteins, for example, given the extensive co-enrichment of both organelles, while the distinctive enrichment of the two organelles in other pull-downs enables a clear differentiation. Overall, proteins from the same compartment share a distinctive enrichment pattern across the whole set of IPs, and can be identified following the principle of correlation profiling^9,10,36^.

Our IP-based strategy extends the collection of methods developed for the proteome-wide characterization of subcellular localization^12,14,16,18,25,26,29,34,49^. An analysis focused on the mitochondrial proteome, for which reference ground-truth is available, revealed a high degree of both precision and recall from our annotations. Furthermore, our dataset enables us to discriminate proteins from closely adjacent compartments. For example, our annotations resolve proteins from the Golgi vs. trans-Golgi, or early endosome vs. recycling endosome, which might be difficult to resolve using centrifugation-based methods because of similar fractionation properties. Spatially adjoined compartments might also be hard to distinguish by proximity ligation; for example outer mitochondrial membrane and peroxisomal proteins cluster together in a recent BioID-based map of the proteome^18^. On the other hand, because organelles are captured whole in a native IP setting, our method cannot directly resolve sub-organellar compartments. For example, our data does not distinguish membrane vs. lumenal proteins, nor different mitochondrial or nuclear sub-compartments, which can be better discriminated by proximity ligation methods^15,18,89^. These examples illustrate how the comprehensive characterization of subcellular architecture will likely require complementary methods applied in parallel.

One advantage of our IP-based workflow is that sample preparation does not require complex instrumentation (e.g., ultracentrifuge), making it relatively simple to implement and scale. Given recent technological development in mass spectrometry (for example the increased sensitivity provided by ion mobility instruments^90^, or data-independent acquisition methods^35^), it is likely that the amount of input material needed for IPs could be significantly reduced from our current scale of 10^7^ cells per replicate. Miniaturizing input amounts would further improve scalability and facilitate applications using cellular models for which large cell cultures might be difficult to obtain, such as cells differentiated from stem cells. IP-based methods also enable the profiling of lipidomes^23^ or metabolomes^21^, paving the way for a characterization of subcellular organization extending beyond proteins.

On top of providing a large data resource, our study also presents a new data-driven, graph-based analytical framework. We show that this framework enables *de novo* identification of subcellular compartments and their inter-connections, and offers an unsupervised formalism to characterize subcellular remodeling. Our approach complements supervised methods that have been developed for the characterization of subcellular localization and dynamics using ground-truth markers as a reference^91^. In addition, we demonstrate that the k-NN representation of our dataset encapsulates, for each protein, specific information that helps defines its function in the cell. Therefore, we envision that our dataset can be used not only as a compendium of annotations of subcellular localization, but also as a resource that can be mined for the generation of specific functional hypotheses.

Finally, we provide a rich characterization of the cellular remodeling induced by infection with the OC43 β-coronavirus. Our data uncovers pervasive subcellular remodeling involving many different cellular compartments and pathways. By capturing responses across the entire cell, our work and others’^32,92^ provide a window to understand the complex strategies that viruses have developed to concurrently hijack a diversity of cellular functions. Strikingly, our results suggest two orthogonal modes of regulation of host proteins, with distinct groups of proteins regulated by changes in their subcellular environment, or by changes in their overall abundance. This underscores the importance of spatial regulation for the control of cellular functions, which is also exemplified by the ubiquity of protein re-localization in signal transduction^7^. Overall, in the same way that spatial methods are transforming transcriptome-based analyses^93^, scalable approaches that can resolve the subcellular coordination of the proteome have the potential to greatly extend our understanding of cellular functions in normal physiology and disease.

## ACKNOWLEDGEMENTS

We sincerely thank R. Zoncu and his team for teaching us native immunoprecipitation protocols. We thank N. Neff and her team for help with high-throughput sequencing; H. Huang, M. Logan, G. Yun, N. Narez, J. Gadiane, and J. Mann, for operational support; and S. Schmid for critical feedback. M.D.L. thanks C. L. Tan for continuous discussions. We thank the Chan Zuckerberg Biohub and its donors Priscilla Chan and Mark Zuckerberg for funding this work.

S. Y.-L. is a Chan Zuckerberg Biohub – San Francisco Investigator.

## SUPPLEMENTARY FIGURE LEGEND

**Supplementary Figure S1 (related to Figure 1).**
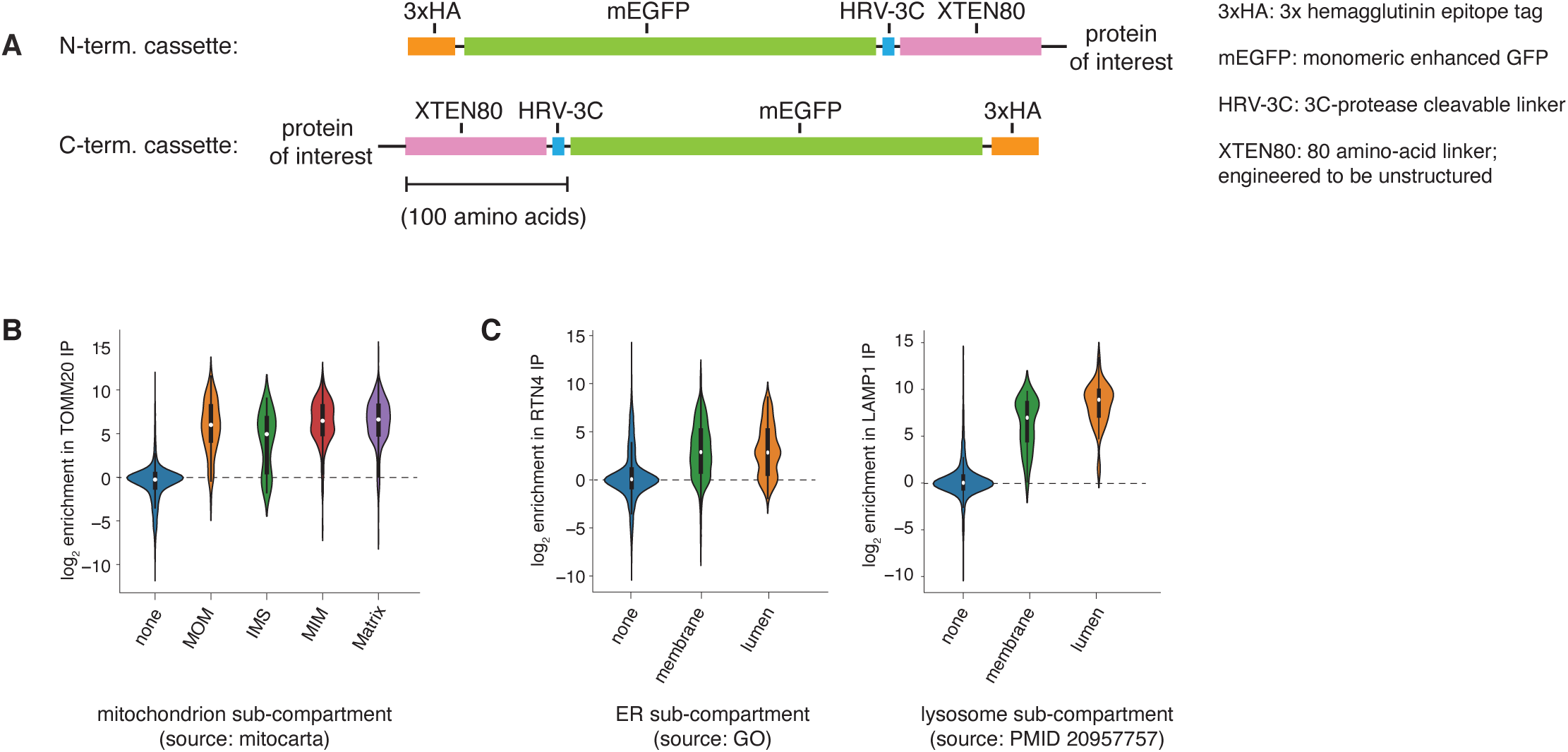
**A)** N- and C-terminal tagging cassettes for organelle immunoprecipitations. **B)** Log2 enrichment of specific categories of mitochondrial proteins in TOMM20 IPs. **C)** Log2 enrichment of membrane vs. lumenal proteins in RTN4 (endoplasmic reticulum) and LAMP1 (lysosome) IPs.

**Supplementary Figure S2 (related to Figure 1).**
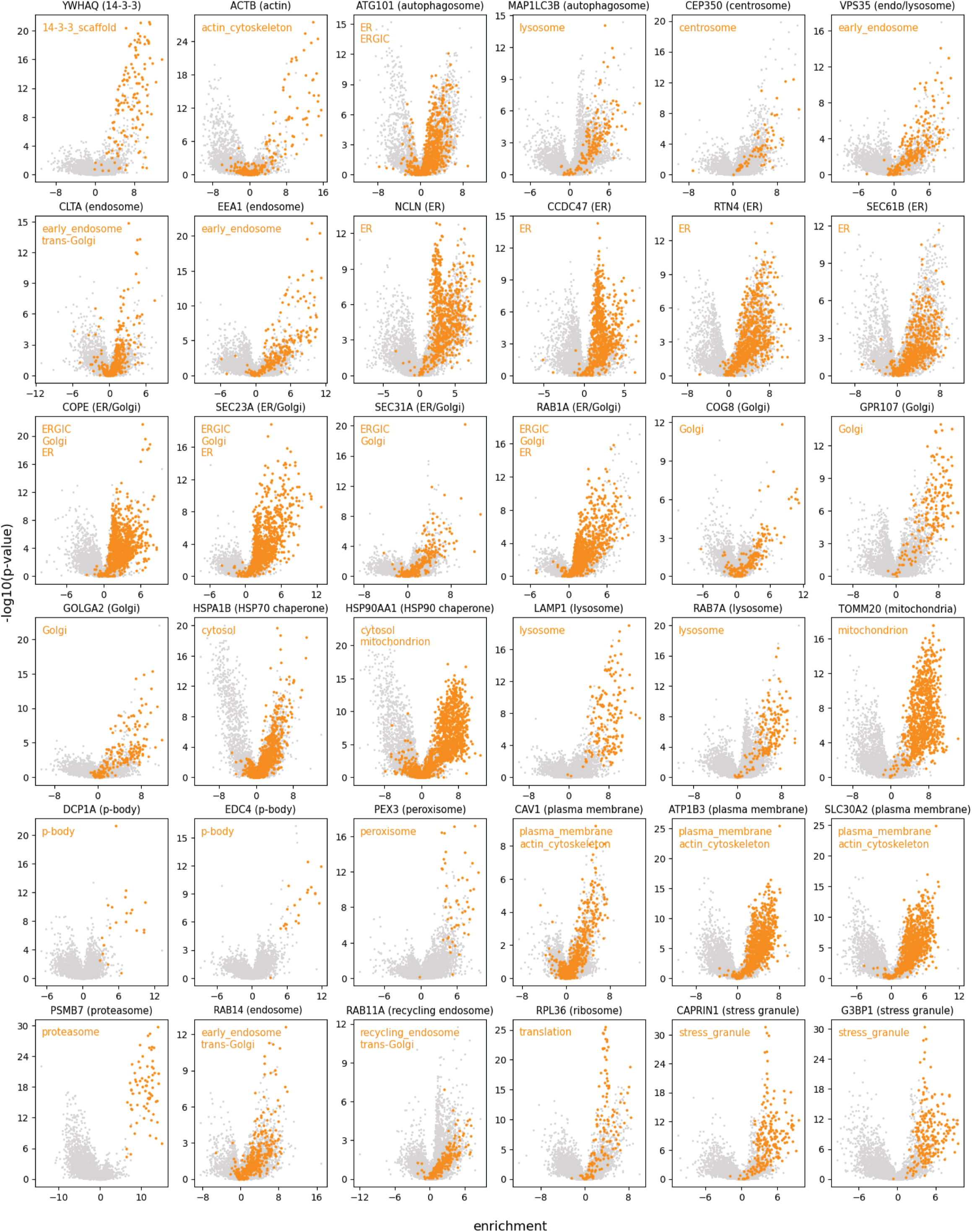
Volcano plots showing protein enrichment in individual IPs from all 36 organelle markers. Plots show log2 enrichment of individual proteins in each IP relative to unrelated controls (See Material and Methods). P-values are calculated from a t-test using triplicate observations of each IP. Proteins annotated to localize to specific sub-cellular compartments are highlighted in orange (annotations from graph-based analysis of our dataset, see text and Suppl. Table 6).

**Supplementary Figure S3 (related to Figure 2).**
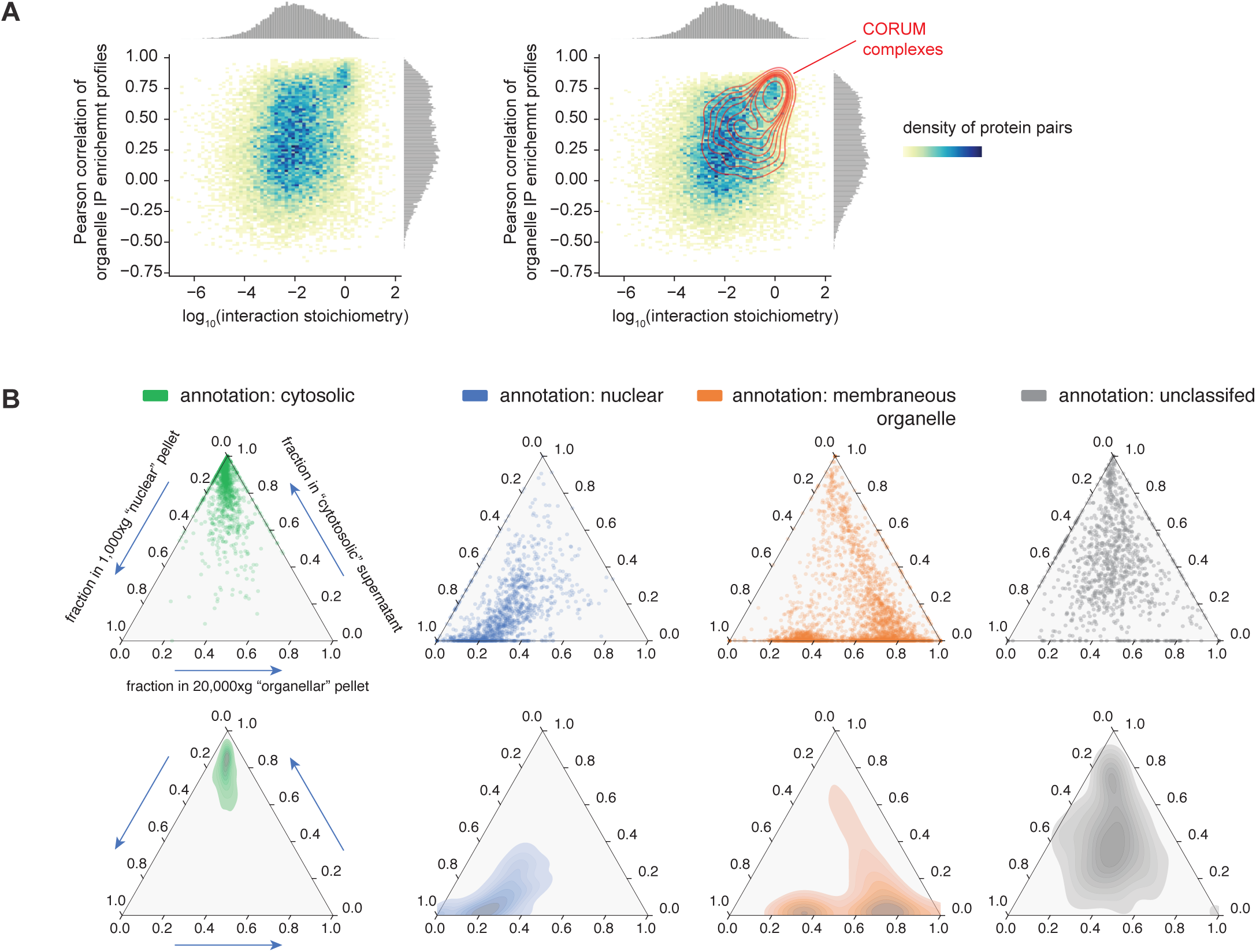
**A)** For each protein pair engaged in direct protein-protein interaction, plots show the relationship between interaction stoichiometry (x-axis, data from the OpenCell database) and similarity of enrichment profile across the whole organelle IP dataset (y-axis, showing Pearson correlation). Both panels display the same data; the right-hand panel highlights stable protein complexes annotated in CORUM. **B)** Ternary plots showing fractional abundance, for individual proteins, in N/O/C spin fractions (cf. Figure 1E). Different groups of proteins are shown in separate panels (from left to right): proteins annotated to localize in the cytosol (green), nucleus (blue), to membranous organelles (orange), and unclassified proteins (grey). While cytosolic, nuclear or organellar proteins show very distinctive enrichment profiles in N/O/C fractions, unclassified proteins do not specifically enrich in any fractions (appearing in the center of the ternary plot), indicating their broad distribution across multiple compartments. Individual proteins are shown as dots in the upper panels, while the lower panels display the overall density of points.

**Supplementary Figure S4 (related to Figure 2).**
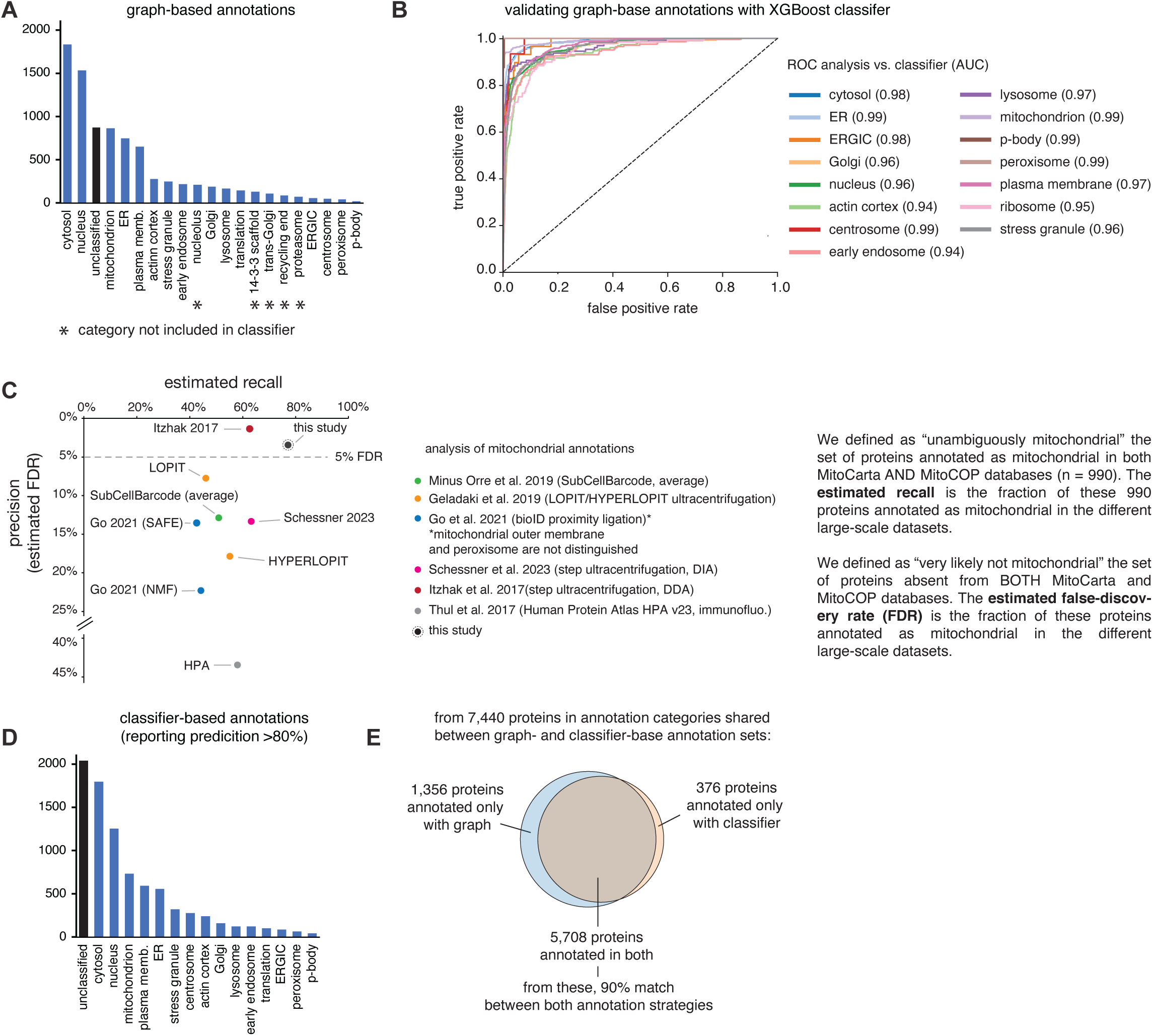
**A)** Protein counts in each graph-based subcellular localization category. **B)** Receiver operating characteristic (ROC) curves of graph-based subcellular localization annotations, using results from an XGBoost classifer as reference “ground-truth”. Corresponding areas under the curve (AUC) values are shown. **C)** Estimation of recall and precision (1 - FDR, false-discovery rate) of mitochondrial annotations in different large-scale datasets. A ground-truth reference of mitochondrial proteins is defined using the MitoCarta and MitoCOP databases (see explanation in the right panel, and text for details). **D)** Protein counts in each classifier-based subcellular localization category. **E)** Comparison of graph-based and classifier-based annotations.

**Supplementary Figure S5 (related to Figure 3).**
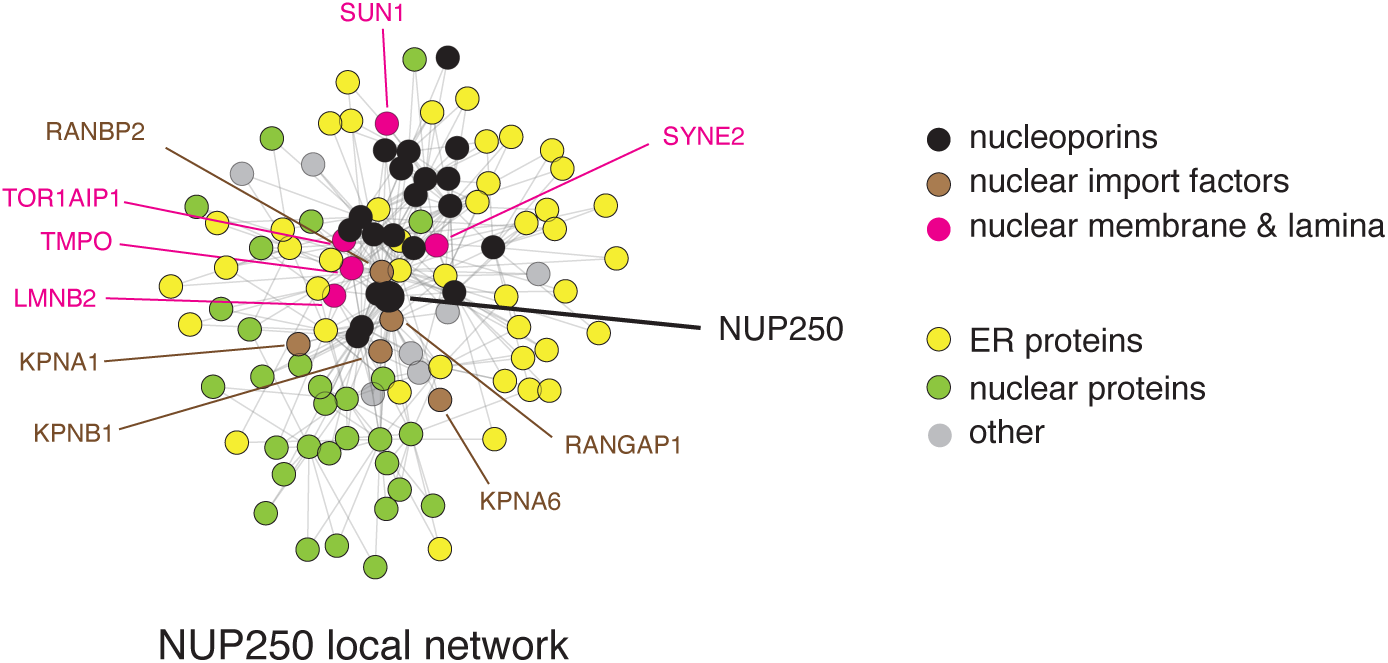
Local k-NN network centered on NUP250, a subunit (nucleoporin) of the nuclear pore complex.

**Supplementary Figure S6 (related to Figure 5).**
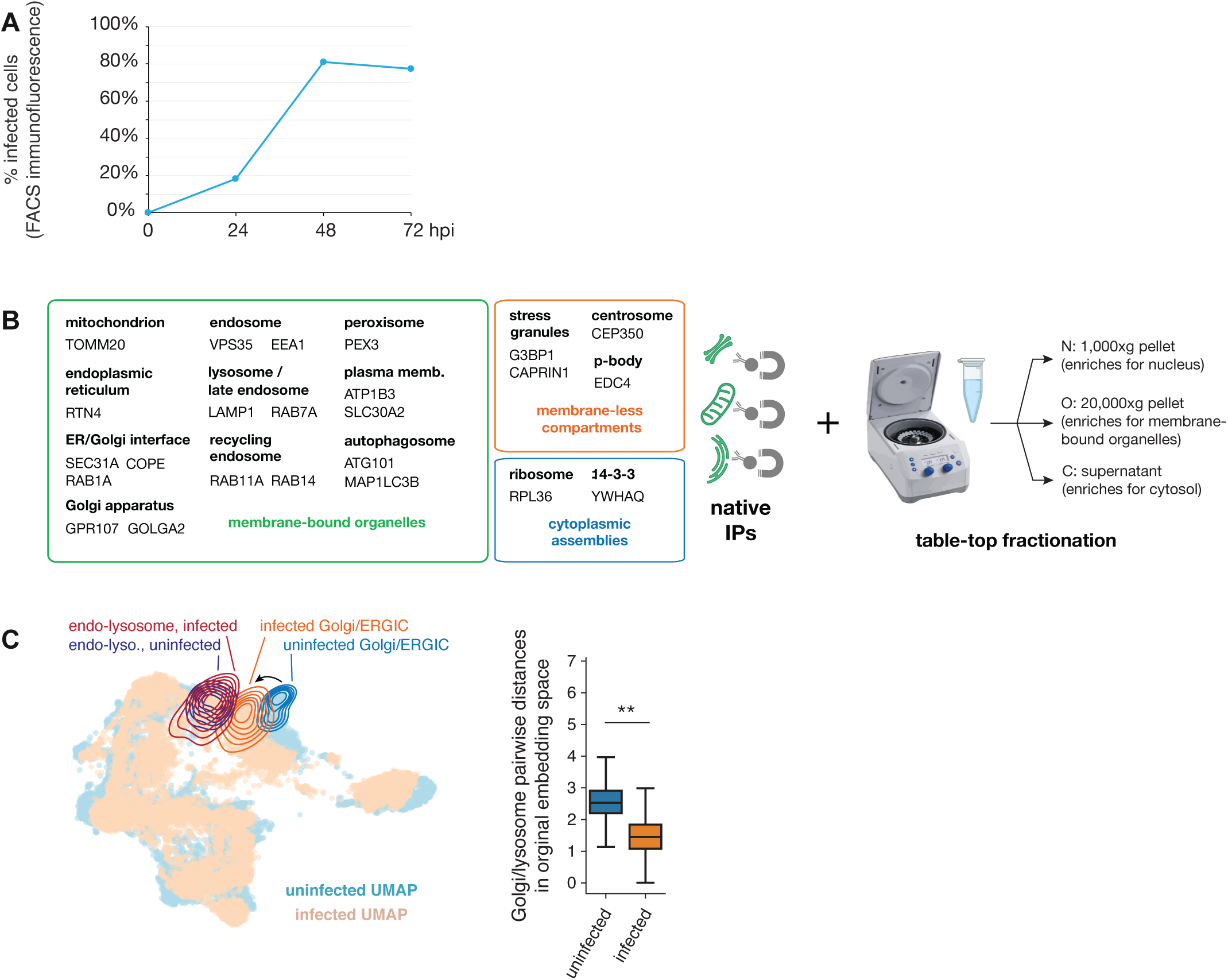
**A)** Percentage of OC43-infected cells as a function of time (hpi: hours post infection). Cells were infected at a MOI of 0.25 at the beginning of the experiment. Infected cells were detected by immunofluorescence flow-cytometry using the monoclonal anti-OC43 antibody 541-8F, exact epitope unknown. **B)** Set of organelle IPs markers included in the OC43 infection experiments. Triplicate IPs were processed side-by-side for uninfected and infected (48 h.p.i, MOI = 0.25) samples for each marker. N/O/C spin fractions were also processed under uninfected and infected conditions. **C)** Distances between Golgi and lysosomal proteins in aligned embedding space are significantly decreased upon infection (p<0.01, t-test). This indicates that the organellar enrichment profiles for Golgi and lysosomal proteins become more similar upon infection.

**Supplementary Figure S7 (related to Figure 6).**
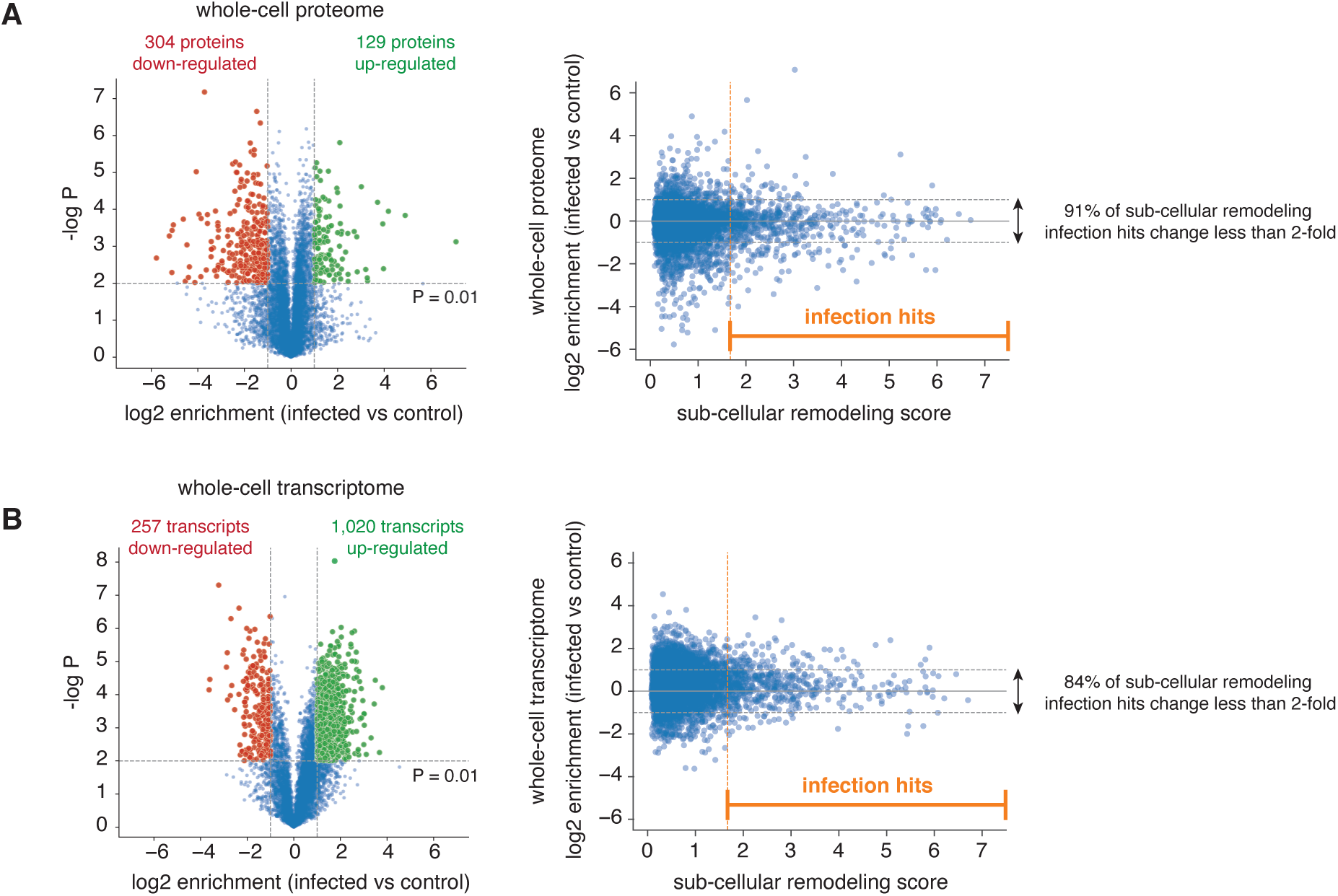
**A)** Whole-cell protein abundance changes upon OC43 infection. Left panel: volcano plot shows log2 enrichment of individual proteins in infected vs. uninfected whole-cell samples. P-values are calculated from a t-test using triplicate observations of each sample. Right panel: log2 enrichment of individual proteins in infected vs. uninfected whole-cell samples is plotted as a function of sub-cellular remodeling scores (cf. Figures 5A-B). **B)** Whole-cell transcript abundance changes upon OC43 infection. Left panel: volcano plot shows log2 enrichment of individual transcripts in infected vs. uninfected whole-cell samples. P-values are calculated from a t-test using triplicate observations of each sample. Right panel: log2 enrichment of individual transcripts and corresponding proteins in infected vs. uninfected whole-cell samples is plotted as a function of sub-cellular remodeling scores (cf. Figures 5A-B).

## MATERIAL AND METHODS

### Cell culture & CRISPR/Cas9 engineering

#### Cell culture

HEK293T cells (ATCC CRL-3216) were cultured in DMEM high-glucose medium (Gibco, cat. #11965118) with 10% fetal bovine serum (Omega Scientific, cat. #FB-11), supplemented with 2mM glutamine (Gibco, cat. #25030081), penicillin and streptomycin (Gibco, cat. #15140163). All cell lines were maintained at 37°C and 5% CO2 and routinely tested for the absence of mycoplasma.

#### Cell line engineering – tagged organelle markers

CRISPR/Cas9 methods were used for gene editing by homology directed repair (HDR) following established protocols. Ribonucleoprotein (RNP) complexes of *S. pyogenes* Cas9 and guide RNA were pre-assembled in vitro, mixed with double-stranded (dsDNA) HDR donors and delivered into HEK293T cells by electroporation in 96-well plates (see below). Each electroporation used 0.2×10^6^ cells, 70 pmol RNP and 2 pmol HDR template. Electroporated cells were incubated for two days in the presence of 1 µM nedisertib (M3814; Selleckchem # S8586) to enhance HDR efficiency^83^. dsDNA HDR donors including tag sequence flanked by gene-specific homology arms were PCR-amplified from a corresponding plasmid template using 5’ biotinylated PCR primers as previously described^84^. RNP complexes were freshly assembled with prior to electroporation, and combined with HDR template in a final volume of 10 µL. First, 0.7 µL gRNA (130 µM stock; source: Integrated DNA technologies) was added to 2.2 µL high-salt RNP buffer {580 mM KCl, 40 mM Tris-HCl pH 7.5, 20% v/v glycerol, 2 mM TCEP-HCl pH 7.5, 2 mM MgCl2, RNAse-free} and incubated at 70°C for 5 min. 1.8 µL of purified Cas9 protein (40 µM stock in Cas9 buffer, ie. 70 pmol) was then added and RNP assembly carried out at 37°C for 10 min. Finally, HDR template (2 pmol) and sterile RNAse-free H2O were added to 10 µL final volume. Electroporation was carried out in Amaxa 96-well shuttle Nucleofector device (Lonza) using SF solution (Lonza) following the manufacturer’s instructions. Cells were washed with PBS and resuspended to 10,000 cells/µL in SF solution (+ supplement) immediately prior to electroporation. For each sample, 20 µL of cells (ie. 200,000 cells) were added to the 10 µL RNP/template mixture. Cells were immediately electroporated using the CM130 program, after which 100 µL of pre-warmed media (containing 1 µM nedisertib) was added to each well of the electroporation plate to facilitate the transfer of 25,000 cells to a new 96-well culture plate containing 150 µL of pre-warmed media (containing 1 µM nedisertib). Electroporated cells were cultured for >5 days and transferred to 12-well plates prior to selection by fluorescence-activated cell sorting (FACS). For each target, 1,200 cells from the top 1% fluorescent cell pool were isolated on a SH800 instrument (Sony biotechnology) and collected in 96-well plates.

#### Cell line engineering – localization de-orphaning

HEK293T cell lines used in the analysis presented in Figure 4 were engineered using the mNeonGreen2(1-10/11) split fluorescent protein system using protocols previously described^14^. In brief, CRISPR/Cas9 methods were used for gene editing by homology directed repair (HDR) using RNP electroporation methods as described in the previous section, with the exception that single-stranded deoxy-oligonucleotide (ssODN) donors were used (Ultramer ssODN, Integrated DNA Technologies; 120 pmol donor per electroporation).

### Sample generation for proteomics analysis

#### Input material

All experiments were performed in triplicate using ∼10^7^ cells per replicate, harvested from 10cm-plate cultures grown to 80-90% confluency.

#### Cell homogenization

Prior to organellar immunoprecipitation or subcellular fractionation, HEK293T cells were mechanically homogenized by repeated passage through a blunt 23G needle in hypotonic buffer. Cells grown on 10cm plates were washed once in ice-cold PBS, and collected by scraping in 5 mL ice-cold PBS. All subsequent steps were conducted at 4 °C or on ice. First, scraped cells were gently pelleted at 500 ×g (4°C) and resuspended in 950 µl of hypotonic homogenization buffer containing: 25 mM Tris-HCl pH 7.5, 50 mM sucrose, 0.2 mM EGTA, 0.5 mM MgCl2, complemented with protease and phosphatase inhibitor cocktail (Thermo Fisher Scientific, #PI78443). Resuspended cells were mechanically homogenized right away by passing the cell suspension 4 times through a 23G blunt 1” needle (SAI Infusion Technology #B23-100) attached to a 1 mL syringe (HSW NormJ-ect-F #NJ-9166017-02), followed by immediate thorough mixing with 89 µL of concentrated sucrose buffer (2.5 M sucrose, 0.2 mM EGTA, 0.5 mM MgCl2) to restore tonicity. Cell homogenates were clarified by centrifugation at 1,000 ×g for 10 min (4°C), resulting in a pellet containing mainly nuclear material, and a supernatant containing mainly organellar and cytosolic proteins.

#### Subcellular fractionation and whole cell extract preparation

For crude subcellular fractionation (“N/O/C” fractions, see text and Figure 1E), the clarified homogenate supernatant was further centrifuged at 20,000 ×g for 45 min (4°C), resulting in a pellet containing mainly organellar proteins, and a soluble cytosolic supernatant. Both the 1,000 ×g (“nucleus”) and 20,000 ×g (“membrane-bound organelles”) pellets were resuspended in SDS lysis buffer {5 % SDS, 50 mM triethylammonium bicarbonate (TEAB) pH 7.55}, 750 µl and 500 µl, respectively. The cytosolic supernatant was supplemented with 20 % SDS to yield 5 % SDS final. All three fractions were boiled for 5 min. In parallel, a whole cell extract (WCE) was generated by directly scraping cells off a 10-cm plate in SDS lysis buffer. All N/O/C and WCE samples were further subjected to probe sonication (UP200St, Hielscher, Germany) and then clarified by centrifugation at 14,000 xg for 15 min. Protein lysates were quantified using BCA assay according to the manufacturer’s instructions (Pierce, Thermo Fisher Scientific). 100 µg of protein was extracted by adding 4.5 volumes of ice-cold acetone, incubated at -20°C for 1 hour, and centrifuged at 21,000 xg at 4°C for 10 min. The protein pellet was then resuspended in {2.5% sodium deoxycholate, 50mM EPPS, pH 8.5} buffer. Protein samples were reduced by adding 1 mM DTT (Thermo Fisher Scientific, A39255) and incubating for 20 min at 37°C. Cysteine side chains were alkylated by adding 5 mM Iodoacetamide (IAA, Thermo Fisher Scientific # A39271) and incubated in the dark for 20 min at room temperature. Samples were digested overnight by adding lys-C (VWR, 100369-826) at an enzyme-to-protein ratio of 1 mAU per 50 µg. Digestion continued the next day by adding trypsin (Thermo Fisher Scientific, 90057) at an enzyme-to-protein ratio of 1:100 for 3 h. All four steps were performed with shaking on a Thermomixer (Eppendorf, Germany). Digestion was stopped by acidification to 1% TFA, incubated on ice for 5 min and centrifuged at 21,000 xg for 15 min to remove insoluble material.

#### Organellar IPs

The input material for organellar IPs is the clarified organellar/cytosolic supernatant (1,000 xg) from the cell homogenization step described above, prepared for cells expressing tagged organellar marker proteins. All steps were carried out at 4°C or on ice. For organelle capture, 750 µL of this supernatant were mixed with 20 µL Anti-HA magnetic beads (Thermo Fisher Scientific #PI88836) in a 96-deepwell plate (1 mL wells), and incubated for 10 min while shaking at 1,000 rpm in a thermomixer. Following capture, beads were bound to a dipping magnet (V&P Scientific #VP 407AM-N1) and washed 3 times in 150 µL ice-cold divalent-free D-PBS (Thermo Fisher Scientific #14190144) by serial transfers to 96-well plates. Beads were released in 150 µL ice-cold D-PBS in a 96-well plate, bound to a flat-bottom magnet (Thermo Fisher Scientific #AM10027), and the D-PBS supernatant removed. Beads were resuspended in 30 µL of DTT-urea buffer (2 M Urea, 12.5 mM Tris-HCl pH 7.5, 1 mM DTT). Bound proteins were alkylated by addition of 3.3 µL of 50 mM Iodoacetamide (IAA, Thermo Fisher Scientific # A39271) to each sample, followed by shaking (1,400 rpm) at RT for 30 min on a thermomixer. Proteins were digested by addition of 0.5 µg of LysC (Fujifilm Wako #NC9223464) and kept shaking overnight at RT. Finally, 1 µg of Trypsin (Thermo Fisher Scientific #PI90058) was added, continuing shaking at RT for 4 h. Beads were removed and the peptides acidified to 1% trifluoroacetic acid (TFA) final.

### Peptide desalting and mass spectrometry

Organellar IP, N/O/C, and WCE peptides were desalted on in-house prepared Styrenedivinylbenzene Reversed Phase Sulfonate packed Stagetips (Rappsilber et al, 2007). Briefly, Stagetips were activated with 100% methanol, conditioned with 80% acetonitrile/0.1% TFA, and equilibrated with 0.2% TFA. Followed by sample loading, washing with 99.9% isopropanol/0.1% TFA, washed twice with 0.2% TFA and once with 0.1% formic acid. Organellar IP peptides were eluted with 60% acetonitrile/0.5% ammonium hydroxide. However, to achieve proteomic depth, N/O/C and WCE peptides were triple fractionated by first eluting with 40% acetonitrile 0.5% TFA 100mM ammonium formate, followed by 60% acetonitrile 0.5% TFA 150mM ammonium formate and finally with 80% acetonitrile 1% ammonium hydroxide. Desalted peptides were flash frozen then dried by centrifugal evaporation and resuspended in 2% acetonitrile/0.1% TFA. Peptides were analyzed on a Fusion Lumos mass spectrometer (Thermo Fisher Scientific, San Jose, CA) equipped with a Thermo EASY-nLC 1200 LC system (Thermo Fisher Scientific, San Jose, CA) and nanoFlex ESI source. Peptides were separated by capillary reverse phase chromatography on a 25 cm column (75 µm inner diameter, packed with 1.6 µm C18 resin, AUR2-25075C18A, Ionopticks, Victoria Australia). Electrospray Ionization voltage was set to 2000 volts. Peptides were introduced into the Fusion Lumos mass spectrometer using a two-step linear gradient. For organellar IP samples, 3–27 % buffer B (0.1% (v/v) formic acid in 80% (v/v) acetonitrile) for 52.5 min followed by 27-40 % buffer B for 14.5 min. For N/O/C and WCE samples, 3–27 % buffer B for 105 min followed by 27-40 % buffer B for 15 min. Both operated with a flow rate of 300 nL/min. Column temperature was maintained at 50°C throughout the procedure. Data was acquired in top speed data-dependent mode with a duty cycle time of 1 s. Full MS scans were acquired in the Orbitrap mass analyzer (FTMS) with a resolution of 120 000 (FWHM) for organellar IP and 240 000 (FWHM) for N/O/C and WCE samples. Both with an m/z scan range of 375-1500 m/z. Selected precursor ions were subjected to fragmentation using higher-energy collisional dissociation (HCD) with quadrupole isolation window of 0.7 m/z, and normalized collision energy of 31%. HCD fragments were analyzed in the Ion Trap mass analyzer (ITMS) set to Turbo scan rate. Fragmented ions were dynamically excluded from further selection for a period of 60 seconds for organellar IP and 45 seconds for N/O/C and WCE samples. The AGC target was set to 1,000,000 and 10,000 for full FTMS and ITMS scans, respectively. The maximum injection time was set to Auto for both full FTMS scans and ITMS scans.

### hCoV-OC43 infection

#### Virus stocks

OC43 was obtained from ATCC (VR-1558) and propagated in Huh7.5.1 cells grown in DMEM media at 34 °C. Viral titers were determined by standard plaque assay using BHK-T7 cells. Briefly, BHK-T7 cells were seeded at 800,000 cells per well in 6-well plates and grown in DMEM media for 24 h at 34°C. The next day, serial 10-fold dilutions of virus stocks were used to infect cells for 2 hours at 34 °C, after which media was removed, and a solution of DMEM media containing 1.2% Avicel RC-591 was overlaid onto the cultures. Infected cells were incubated for 6 days at 34 °C, followed by fixation with 4 % formaldehyde, staining with crystal violet, and plaque counting. All experiments were performed in a biosafety level 2 laboratory.

#### Infection – organelle IPs

For organelle IPs from infected cells, HEK293T cells expressing tagged organellar markers were seeded ∼18 h before infection at a density of ∼5*10^6^ per 10 cm plate, resulting in a confluency of ∼80 % at the time of infection. Infection was carried out by direct addition of viral inoculum into the media, at a multiplicity of infection (MOI) of 0.25, at 37 °C. This MOI was optimized to result in a uniformly infected population at 48 h.p.i. while minimizing the required amount of inoculum. Cells were then harvested and processed like uninfected cells described above.

#### Infection – imaging

The day prior to infection, cells were seeded on 96-well glass-bottom plates (Greiner Bio One, cat. #655891) coated with 50 µg/mL fibronectin (Corning, cat. #356008; manufacturer’s protocol). 4000-8000 cells per well were seeded for uninfected conditions, and at least 8000 cells per well were seeded for infected conditions. Day of infection, virus aliquots were thawed on ice, diluted in growth media, and applied onto cells at indicated MOI and at a final volume of 100 µL/well. For MOI calculations, cells were assumed to have tripled. MOI 0 conditions simply received media change. Unless indicated otherwise, infection imaging experiments were live-imaged at 48 hpi. In infection experiments co-treated with Ferrostatin (Selleck Chemicals, cat. # S7243), cells were first treated with 50 µL/well of virus at the desired MOI, and then immediately treated with 50 µL/well of 2x desired Ferrostatin concentration to reach a final volume of 100 µL/well.

### Microscopy imaging

#### Sample preparation

Live-cell imaging was performed on 96-well glass-bottom plates (Greiner Bio One, cat. #655891) coated with 50µg/ml fibronectin (Corning, cat. #356008). For cells to be imaged the next day, cells were seeded on an imaging plate ∼30 hours before imaging at 15,000 cells per well. Before imaging, cells were counter-stained with the live-cell DNA dye Hoechst 33342 (Invitrogen, cat. #H3570) by incubation for 30 minutes at 37°C in 150 µl of Hoechst diluted to 1µg/mL in culture media. Media was then replaced with phenol-free DMEM (Gibco, cat. #21063029) supplemented with 10% FBS. Hoechst staining was performed three to four hours prior to imaging to provide the cells time to recover from any mechanical stress due to medium changes. In experiments involving FeRhoNox-1 (Sigma-Aldrich, cat. #SCT030), FeRhoNox-1 was resuspended following manufacturer’s protocol and used at a final concentration of 5 µM in combination with Hoechst. Cells were then incubated at 37°C for 1.5 hours in serum-free imaging media, and subsequently washed and imaged in serum-free imaging media.

#### Live-cell fluorescence microscopy

Cells were imaged on a DMI-8 inverted microscope (Leica) equipped with a Dragonfly spinning-disk confocal system (Andor), a 63x 1.47NA oil objective (Leica), and a 16-bit iXon Ultra 888 EMCCD camera (Andor, pixel size: 13×13 µm2). A pinhole size of 40μm was used with an EM gain of 400. Cells were maintained at 37 °C and 5% CO2 during image acquisition by a stage-top incubator (Okolab, H101-K-Frame). The microscope was controlled using the open-source microscope-control software MicroManager (version 1.4.22).

### Flow cytometry

70000 cells per well were seeded into 12 well plastic plates day prior to infection. At 48 h post MOI 1 infection (final volume of 1 mL/well), cells were resuspended in trypsin, quenched, and fixed in 4% paraformaldehyde at room temperature for 15 min. In experiments involving FeRhoNox-1 (see “Sample Preparation” section for protocol), cells were treated with FeRhoNox-1 prior to fixation. Cells were then washed, permeabilized, and blocked using Perm/Wash buffer (BD Biosciences, cat. #554723; >30 min at room temperature). For antibody staining, cells were treated with 1:200 OC43 antibody (Millipore Sigma, cat. #MAB9012; >1 h at room temperature), washed, treated with 1:500 Alexa Fluor secondary antibody (Thermo Fisher Scientific; >30 min at room temperature), and washed (all steps here in Perm/Wash buffer). Finally, cells were resuspended in FACS buffer (HBSS with no ions (Gibco, cat. #14175095) with 1% FBS (Omega Scientific, cat. #FB-11) and 25 mM HEPES (Gibco, cat. #15630080)) and analyzed using CytoFLEX (Beckman Coulter).

### Transcriptomics

For transcriptomics of OC43-infected cells, we cultured and infected cells essentially as for all proteomics experiments, on 10 cm plates at 37 °C. Cells were infected at ∼80 % confluency with an MOI of 1.0. We staggered a series of infections so that samples were harvested in parallel in an uninfected state and at 16, 24, and 48 h.p.i, respectively. Cells were harvested by trypsinization, quenched, and washed in PBS. We then used MULTI-seq^94^ to encode the experimental conditions (i.e. uninfected or 16, 24, 48 h.p.i) before pooling all cells and loading them onto one lane of a 10x chip (NextGEM Single-Cell V(D)J Reagents Kit v1.1, 10x Genomics), aiming for recovery of 20,000 cells. We used the 5′ workflow to distinguish the different viral subgenomic RNAs^95^. We followed the manufacturer’s recommendation for library preparation and sequencing, spiking in the MULTI-seq amplicons at ∼8 % of the gene expression library, at ∼50,000 reads per cell. We processed the raw data with cellranger v. 5.0.1 using a custom transcriptome reference that combines the GRCh38 human genome and a manually curated reference for OC43. In addition to the default cellranger filtering, we removed cells with <2,000 UMIs, cells with >5% content of mitochondrial transcripts, as well as cells whose MULTI-seq barcodes indicated multiplets or could not be unambiguously assigned to one experimental condition, resulting in 12,910 cells in the final dataset. We used leiden clustering to define three main cellular states, corresponding to uninfected cells, and cells with low and high viral loads, respectively. To assess differential gene expression between uninfected and infected cells presenting high viral load, we generated pseudo-bulk transcriptomes from sets of 100 randomly subsampled cells from either cell state.

### Data analysis – mass spectrometry proteomics

#### Data availibity

Mass spectrometry raw data and associated MaxQuant output tables were deposited to the ProteomeXchange Consortium via the PRIDEpartner repository (accession PXD046440).

#### Raw Data Processing

Raw data were processed in MaxQuant^96^ v. 2.2.0.0. We defined separate parameter groups for organellar IPs and used the match between runs feature and MaxLFQ^97^ for label-free quantification (using 1 min. LFQ ratio count), each within those groups. Spectra were searched against human protein sequences (downloaded from Uniprot on 2021-07-17, canonical and isoform sequences) and 23 manually compiled hCoV-OC43 protein sequences (assembled based on the genome of hCoV-OC43 strain 1588, provided by ATCC). Briefly, a 1% false discovery rate was set at the peptide spectrum and protein matching with a minimum peptide length of 7, 2 missed cleavages were allowed, and enzyme was set to trypsin. Oxidation of methionine and protein N-terminal acetylation were set as variable modifications along with carbamidomethylation of cysteines set as a fixed modification.

#### Protein identification in native immunoprecipitation

Protein identifications were filtered, removing common contaminants, hits to the reverse decoy database as well as proteins only identified through post-translational modifications at specific sites. We used log2 MaxQuant LFQ intensities for all analyses. We required that each protein be quantified in all replicates from at least one native immunoprecipitation-mass-spectrometry (IP-MS) sample and in two replicates of whole proteome MS samples. Missing values were addressed by imputation using a Gaussian distribution, centered on the left tail of the global distribution of LFQ intensities. Specifically, the offset of the mean of this Gaussian distribution is 1.8 times the global standard deviation, while its standard deviation is fixed at 0.3 times the global standard deviation.

#### Protein enrichment analysis using affinity-enrichment mass spectrometry

Our strategy for enrichment analysis uses the established concept of affinity-enrichment mass spectrometry (AE-MS)^85^. In AE-MS, rather than using a single negative control to measure protein enrichment profiles for a given IP-MS sample, enrichment is calculated relative to a larger set of unrelated IP-MS samples (that is, a larger set of samples that do not specifically capture the same set of proteins as the sample of interest). As previously described^14,85,86^, this enables a better estimation of the null distribution of background binders and leads to a more robust identification of interactions.

#### Sample grouping

For the current study, the full set of triplicate IP-MS samples was generated across 17 separate experimental batches, each batch containing on average 8 triplicate samples (or 24 individual IPs). For each batch, we first defined the background of proteins that bind or adsorb to the IP beads in a manner that is not 3xHA tag-specific; this non-specific background is defined by the LFQ intensities measured in a triplicate set of negative control IP-MS included in each batch, obtained using wild-type (untagged) cells. To minimize batch effects, we then grouped together batches that shared similar background intensity profiles using hierarchical clustering of these negative controls (see details on Github). This analysis defined 5 overall groups of samples.

#### Enrichment calculation

All enrichment calculations use triplicate sets of measurements, which will be subsequently described as “samples”. Within each of the five sample groups, raw protein enrichment values were first computed using the cohort-matching, untagged negative control as the null distribution. The pairwise correlation of raw protein enrichment between all samples in a group was calculated, and uncorrelated samples were identified (correlation coefficient <0.35). Final protein enrichment values were then calculated using, for each sample, the full set of other uncorrelated samples from the same group to define a null distribution. The final enrichment value for each protein was calculated by taking the difference in median log2 LFQ values between the replicates and the null distribution. The significance of the final enrichment was tested using a t-test comparing the mean of log2 LFQ values between the replicates and the null distribution.

#### Calculation of protein proportions in N/O/C spin-fractions

Protein identifications in spin-fraction MS samples were filtered using identical methods outlined for IP-MS samples. We required that each protein be quantified in at least one out of three spin-fractions. The N/O/C proportions for each protein (from triplicate experiments) were defined by dividing median N, O or C LFQ values by the sum of median LFQ values obtained from all three spin-fractions.

### Graph-based analyses

#### k-nearest neighbor (k-NN) graph

To construct the k-NN graph, we combined the protein enrichment data matrix with the N/O/C proportions data matrix. This integration required the presence of the same proteins in both matrices (that is, proteins absent from one of the matrices were excluded form analysis). We employed the *scanpy* package^98^ for the construction of the k-NN graph. In this process, the combined data matrix was first scaled to achieve zero mean and unit variance with clipping values at 10. Subsequently, we applied the UMAP algorithm^41^, using 20 nearest neighbors and the Euclidean distance metric, to generate a weighted adjacency matrix delineating the edges of the k-NN graph.

#### Dimension reduction

The combined matrix of enrichment value and N/O/C proportions were scaled to achieve zero mean and unit variance with clipping values at 10. The UMAP algorithm^41^ was used to embed the scaled matrix in two or three-dimensional space using 20 nearest neighbors, the Euclidean distance metric, and a minimum embedding distance of 0.1.

#### Clustering and protein-level compartment annotation

The clustering of protein groups was computed with the Leiden algorithm implemented in the *scanpy* package^98^. The k-NN graph was used as input and the “partition_type” parameter was set to “leidenalg.RBConfigurationVertexPartition”. Protein-level compartment annotations were obtained through ontology enrichment analysis. This analysis was conducted using the Enrichr^99^ API at https://maayanlab.cloud/speedrichr. For each Leiden cluster, enrichment was calculated against a background list inlcuding all proteins detected in our dataset. Enrichment analysis was computed using the COMPARTMENTS^44^ and GO-Cellular component^45^ databases using a p-value cutoff of 0.01. The most significantly enriched gene ontology term for each Leiden cluster was then assigned as the annotation for the proteins in that cluster. Leiden clusters with the same annotation were merged.

#### Graph-based annotation

Protein-level compartment annotations were refined using the k-NN graph. For each protein, the compartment annotation most frequently observed among all its neighbors in the graph was adopted as the protein’s graph-based annotation. This approach excluded the “unclassified” category from being selected as the most frequent annotation.

#### Jaccard coefficient

To calculate a Jaccard coefficient, two partites have to be defined to which a given node might be connected to. Therefore, for a given protein, we identified which two compartment annotations (partites) were the most found in the list of that protein’s direct neighbors in the k-NN graph. Compartments comprising large numbers of proteins overall might have an advantage in this calculation. Therefore, to rank the top-two partite compartments for each protein while accounting for the size of different compartments, the number of that protein’s neighbors annotated as belonging to a given compartment was normalized by the total size of that compartment. A minimum threshold was also established: to rank as one of the top-two partites, a compartment must contain three or more of the direct k-NN neighbors of given protein. Jaccard coefficients were computed by dividing the sum of neighbors in the two selected compartments by the sum of the size of the two compartments subtracting the sum of neighbors. Cluster connectivity. Connectivity between any two given graph-based annotation clusters in the k-NN graph was quantified by calculating the percentage of connections between two clusters over the total possible connections that could exist between them.

### Classifier-based analysis

The compartment classifier was built using the eXtreme Gradient Boosting (XGBoost) algorithm^47^. implemented via the *XGBoost* package^100^. The training dataset comprises the combined matrix of enrichment values and N/O/C proportions. In the preprocessing stage, this matrix was scaled to achieve zero mean and unit variance with clipping values at 10. The training dataset comprised 61 IP-MS/spin-fraction-MS samples as features and 2088 proteins as observation instances. Out of these 2088 proteins, 1288 were from 13 organellar compartments with substantial reference evidence (Suppl. Table 1), supplemented by 400 cytosolic and 400 nuclear proteins. These latter proteins were selectively subsampled from high confidence predictions in Itzhak et al.^34^ to avoid introducing heavily overrepresented classes. To further address class imbalance, we employed a hybrid resampling technique known as SMOTENN^101^, which combines both over- and under-sampling.

Hyperparameter tuning of the XGBoost model was conducted using the Bayesian optimization algorithm^102^, implemented via the *scikit-optimize* package (https://zenodo.org/records/5565057). The optimization process tuned the following parameters: “max_depth”, “learning_rate”, “gamma”, and “grow_policy”, using “f1_weighted” as the primary scoring metric. Three-fold cross-validation was applied in all model training processes (including hyperparameter tuning) to minimize the effect of randomly splitting training data into train and test subsets. During training, the model performance was monitored using “mlogloss”, “merror”, and “auc” metrics computed from the test subset. An “early_stopping_rounds” of 50 was configured to prevent model overfitting. The final evaluation of the model was performed with 5183 proteins that are not in the training dataset.

Classifier-based annotations were derived utilizing an XGBoost model trained using a set of optimized hyperparameters (see Github for details). The “objective” parameter was set to “multi:softprob” to allow the generation of probabilistic predictions for each compartment. For each group of proteins, the compartment associated with the highest prediction probability was identified and designated as the classifier-based annotation.

### Subcellular remodeling analysis

#### Aligned-UMAP

Protein identifications from IP-MS and N/O/C spin-fractions MS samples were separately processed for uninfected control and infected conditions following identical methods outlined above. As a result, two combined matrices were generated: one representing the uninfected control condition and the other for the infected condition. These matrices were then utilized to construct respective k-NN graphs. Both graphs were simultaneously embedded into a shared low-dimensional space in an aligned manner using the AlignedUMAP method, as implemented in the umap package ( see <u>umap-learn.readthedocs.io/en/latest/</u> <u>aligned_umap_basic_usage.html</u>). The embedding was configured with the following parameters: 20 nearest neighbors, Euclidean distance metric, minimum embedding distance 0.1, 300 epochs, and an alignment regularization factor of 0.002.

#### Subcellular remodeling score

The uninfected and infected k-NN graphs were embedded into a shared 10-dimensional space using the AlignedUMAP method described above. To minimize stochasticity, we repeated the embedding process for 200 times. For each protein group, the remodeling score is defined as the average Euclidean distance between the infected and uninfected coordinates of 200 repetitions.

#### Visualization of protein subcellular remodeling

The uninfected and infected k-NN graphs were embedded into a shared 2-dimensional space for comparative visual representation. Protein-level compartment annotations were generated using the methodology outlined above, with a notable modification: the resolution parameter in the Leiden algorithm was adjusted to a value of 1.

### Figure generation and general code availability

Data analysis for this study was conducted using the Python programming language. Figures were generated in Python using matplotlib, seaborn or plotly packages. All computer code and reference data used to conduct the analysis and generate the figures in this manuscript can be found on GitHub at <u>github.com/czbiohub-sf/Organelle_IP_analyses_and_figures</u>.

## BIBLIOGRAPHY

1. Carroll, M. (1989). Organelles. 1–18. 10.1007/978-1-349-19781-1_1.

2. Cohen, S., Valm, A.M., and Lippincott-Schwartz, J. (2018). Interacting organelles. Curr. Opin. Cell Biol. 53, 84–91. 10.1016/j.ceb.2018.06.003.

3. Wheeler, R.J., and Hyman, A.A. (2018). Controlling compartmentalization by non-membrane-bound organelles. Philos. Trans. R. Soc. B: Biol. Sci. 373, 20170193. 10.1098/rstb.2017.0193.

4. Hirose, T., Ninomiya, K., Nakagawa, S., and Yamazaki, T. (2023). A guide to membraneless organelles and their various roles in gene regulation. Nat. Rev. Mol. Cell Biol. 24, 288–304. 10.1038/s41580-022-00558-8.

5. Bar-Peled, L., and Kory, N. (2022). Principles and functions of metabolic compartmentalization. Nat. Metab. 4, 1232–1244. 10.1038/s42255-022-00645-2.

6. Speijer, D., Manjeri, G.R., and Szklarczyk, R. (2014). How to deal with oxygen radicals stemming from mitochondrial fatty acid oxidation. Philos. Trans. R. Soc. B: Biol. Sci. 369, 20130446. 10.1098/rstb.2013.0446.

7. Bauer, N.C., Doetsch, P.W., and Corbett, A.H. (2015). Mechanisms Regulating Protein Localization. Traffic 16, 1039–1061. 10.1111/tra.12310.

8. Schlacht, A., Herman, E.K., Klute, M.J., Field, M.C., and Dacks, J.B. (2014). Missing Pieces of an Ancient Puzzle: Evolution of the Eukaryotic Membrane-Trafficking System. Cold Spring Harb. Perspect. Biol. 6, a016048. 10.1101/cshperspect.a016048.

9. Lundberg, E., and Borner, G.H.H. (2019). Spatial proteomics: a powerful discovery tool for cell biology. Nature Reviews Molecular Cell Biology 20, 285–302. 10.1038/s41580-018-0094-y.

10. Christopher, J.A., Stadler, C., Martin, C.E., Morgenstern, M., Pan, Y., Betsinger, C.N., Rattray, D.G., Mahdessian, D., Gingras, A.-C., Warscheid, B., et al. (2021). Subcellular proteomics. Nat. Rev. Methods Primers 1, 32. 10.1038/s43586-021-00029-y.

11. Christopher, J.A., Geladaki, A., Dawson, C.S., Vennard, O.L., and Lilley, K.S. (2022). Subcellular Transcriptomics and Proteomics: A Comparative Methods Review. Mol Cell Proteomics 21, 100186. 10.1016/j.mcpro.2021.100186.

12. Thul, P.J., Åkesson, L., Wiking, M., Mahdessian, D., Geladaki, A., Blal, H.A., Alm, T., Asplund, A., Björk, L., Breckels, L.M., et al. (2017). A subcellular map of the human proteome. Science 356, eaal3321. 10.1126/science.aal3321.

13. Huh, W.-K., Falvo, J.V., Gerke, L.C., Carroll, A.S., Howson, R.W., Weissman, J.S., and O’Shea, E.K. (2003). Global analysis of protein localization in budding yeast. Nature 425, 686– 691. 10.1038/nature02026.

14. Cho, N.H., Cheveralls, K.C., Brunner, A.-D., Kim, K., Michaelis, A.C., Raghavan, P., Kobayashi, H., Savy, L., Li, J.Y., Canaj, H., et al. (2022). OpenCell: Endogenous tagging for the cartography of human cellular organization. Science 375, eabi6983. 10.1126/science.abi6983.

15. Rhee, H.-W., Zou, P., Udeshi, N.D., Martell, J.D., Mootha, V.K., Carr, S.A., and Ting, A.Y. (2013). Proteomic mapping of mitochondria in living cells via spatially restricted enzymatic tagging. Science (New York, N.Y.) 339, 1328–1331. 10.1126/science.1230593.

16. Liu, X., Salokas, K., Tamene, F., Jiu, Y., Weldatsadik, R.G., Öhman, T., and Varjosalo, M. (2018). An AP-MS- and BioID-compatible MAC-tag enables comprehensive mapping of protein interactions and subcellular localizations. Nat. Commun. 9, 1188. 10.1038/s41467-018-03523-2.

17. Youn, J.-Y., Dunham, W.H., Hong, S.J., Knight, J.D.R., Bashkurov, M., Chen, G.I., Bagci, H., Rathod, B., MacLeod, G., Eng, S.W.M., et al. (2018). High-Density Proximity Mapping Reveals the Subcellular Organization of mRNA-Associated Granules and Bodies. Mol Cell 69, 517–532.e11. 10.1016/j.molcel.2017.12.020.

18. Go, C.D., Knight, J.D.R., Rajasekharan, A., Rathod, B., Hesketh, G.G., Abe, K.T., Youn, J.-Y., Samavarchi-Tehrani, P., Zhang, H., Zhu, L.Y., et al. (2021). A proximity-dependent biotinylation map of a human cell. Nature, 1–5. 10.1038/s41586-021-03592-2.

19. Chen, W.W., Freinkman, E., Wang, T., Birsoy, K., and Sabatini, D.M. (2016). Absolute Quantification of Matrix Metabolites Reveals the Dynamics of Mitochondrial Metabolism. Cell 166, 1324–1337.e11. 10.1016/j.cell.2016.07.040.

20. Chen, W.W., Freinkman, E., and Sabatini, D.M. (2017). Rapid immunopurification of mitochondria for metabolite profiling and absolute quantification of matrix metabolites. Nature protocols 12, 2215–2231. 10.1038/nprot.2017.104.

21. Abu-Remaileh, M., Wyant, G.A., Kim, C., Laqtom, N.N., Abbasi, M., Chan, S.H., Freinkman, E., and Sabatini, D.M. (2017). Lysosomal metabolomics reveals V-ATPase- and mTOR-dependent regulation of amino acid efflux from lysosomes. Science (New York, N.Y.) 358, 807– 813. 10.1126/science.aan6298.

22. Ray, G.J., Boydston, E.A., Shortt, E., Wyant, G.A., Lourido, S., Chen, W.W., and Sabatini, C. M. (2020). A PEROXO-Tag Enables Rapid Isolation of Peroxisomes from Human Cells. iScience 23. 10.1016/j.isci.2020.101109.

23. Park, H., Hundley, F.V., Yu, Q., Overmyer, K.A., Brademan, D.R., Serrano, L., Paulo, J.A., Paoli, J.C., Swarup, S., Coon, J.J., et al. (2022). Spatial snapshots of amyloid precursor protein intramembrane processing via early endosome proteomics. Nat. Commun. 13, 6112. 10.1038/s41467-022-33881-x.

24. Fasimoye, R., Dong, W., Nirujogi, R.S., Rawat, E.S., Iguchi, M., Nyame, K., Phung, T.K., Bagnoli, E., Prescott, A.R., Alessi, D.R., et al. (2023). Golgi-IP, a tool for multimodal analysis of Golgi molecular content. Proc National Acad Sci 120, e2219953120. 10.1073/pnas.2219953120.

25. Orre, L.M., Vesterlund, M., Pan, Y., Arslan, T., Zhu, Y., Woodbridge, A.F., Frings, O., Fredlund, E., and Lehtiö, J. (2019). SubCellBarCode: Proteome-wide Mapping of Protein Localization and Relocalization. Mol Cell 73, 166–182.e7. 10.1016/j.molcel.2018.11.035.

26. Martinez-Val, A., Bekker-Jensen, D.B., Steigerwald, S., Koenig, C., Østergaard, O., Mehta, A., Tran, T., Sikorski, K., Torres-Vega, E., Kwasniewicz, E., et al. (2021). Spatial-proteomics reveals phospho-signaling dynamics at subcellular resolution. Nat. Commun. 12, 7113. 10.1038/s41467-021-27398-y.

27. Morgenstern, M., Peikert, C.D., Lübbert, P., Suppanz, I., Klemm, C., Alka, O., Steiert, C., Naumenko, N., Schendzielorz, A., Melchionda, L., et al. (2021). Quantitative high-confidence human mitochondrial proteome and its dynamics in cellular context. Cell Metab 33, 2464–2483.e18. 10.1016/j.cmet.2021.11.001.

28. Andersen, J.S., Wilkinson, C.J., Mayor, T., Mortensen, P., Nigg, E.A., and Mann, M. (2003). Proteomic characterization of the human centrosome by protein correlation profiling. Nature 426, 570–574. 10.1038/nature02166.

29. Foster, L.J., Hoog, C.L. de, Zhang, Y., Zhang, Y., Xie, X., Mootha, V.K., and Mann, M. (2006). A Mammalian Organelle Map by Protein Correlation Profiling. Cell 125, 187–199. 10.1016/j.cell.2006.03.022.

30. Dunkley, T.P.J., Hester, S., Shadforth, I.P., Runions, J., Weimar, T., Hanton, S.L., Griffin, J.L., Bessant, C., Brandizzi, F., Hawes, C., et al. (2006). Mapping the Arabidopsis organelle proteome. Proc. Natl. Acad. Sci. 103, 6518–6523. 10.1073/pnas.0506958103.

31. Borner, G.H.H., Hein, M.Y., Hirst, J., Edgar, J.R., Mann, M., and Robinson, M.S. (2014). Fractionation profiling: a fast and versatile approach for mapping vesicle proteomes and protein– protein interactions. Molecular Biology of the Cell 25, 3178–3194. 10.1091/mbc.e14-07-1198.

32. Jean Beltran, P.M., Mathias, R.A., and Cristea, I.M. (2016). A Portrait of the Human Organelle Proteome In Space and Time during Cytomegalovirus Infection. Cell Systems 3, 361–373.e6. 10.1016/j.cels.2016.08.012.

33. Christoforou, A., Mulvey, C.M., Breckels, L.M., Geladaki, A., Hurrell, T., Hayward, P.C., Naake, T., Gatto, L., Viner, R., Arias, A.M., et al. (2016). A draft map of the mouse pluripotent stem cell spatial proteome. Nat Commun 7, 8992. 10.1038/ncomms9992.

34. Itzhak, D.N., Tyanova, S., Cox, J., and Borner, G.H. (2016). Global, quantitative and dynamic mapping of protein subcellular localization. Cell 5, 570. 10.7554/elife.16950.

35. Schessner, J.P., Albrecht, V., Davies, A.K., Sinitcyn, P., and Borner, G.H.H. (2023). Deep and fast label-free Dynamic Organellar Mapping. Nat. Commun. 14, 5252. 10.1038/s41467-023-41000-7.

36. Borner, G.H.H. (2020). Organellar Maps Through Proteomic Profiling – A Conceptual Guide. Mol. Cell. Proteom. 19, 1076–1087. 10.1074/mcp.r120.001971.

37. Fransen, M., Lismont, C., and Walton, P. (2017). The Peroxisome-Mitochondria Connection: How and Why? Int. J. Mol. Sci. 18, 1126. 10.3390/ijms18061126.

38. Rowland, A.A., and Voeltz, G.K. (2012). Endoplasmic reticulum–mitochondria contacts: function of the junction. Nat. Rev. Mol. Cell Biol. 13, 607–615. 10.1038/nrm3440.

39. Decker, C.J., and Parker, R. (2012). P-Bodies and Stress Granules: Possible Roles in the Control of Translation and mRNA Degradation. Cold Spring Harb. Perspect. Biol. 4, a012286. 10.1101/cshperspect.a012286.

40. Giurgiu, M., Reinhard, J., Brauner, B., Dunger-Kaltenbach, I., Fobo, G., Frishman, G., Montrone, C., and Ruepp, A. (2018). CORUM: the comprehensive resource of mammalian protein complexes—2019. Nucleic Acids Res 47, gky973-. 10.1093/nar/gky973.

41. McInnes, L., Healy, J., and Melville, J. (2018). UMAP: Uniform Manifold Approximation and Projection for Dimension Reduction. Arxiv.

42. Traag, V.A., Waltman, L., and Eck, N.J. van (2019). From Louvain to Leiden: guaranteeing well-connected communities. Sci Rep-uk 9, 5233. 10.1038/s41598-019-41695-z.

43. Wolf, F.A., Hamey, F.K., Plass, M., Solana, J., Dahlin, J.S., Göttgens, B., Rajewsky, N., Simon, L., and Theis, F.J. (2019). PAGA: graph abstraction reconciles clustering with trajectory inference through a topology preserving map of single cells. Genome Biol. 20, 59. 10.1186/s13059-019-1663-x.

44. Binder, J.X., Pletscher-Frankild, S., Tsafou, K., Stolte, C., O’Donoghue, S.I., Schneider, R., and Jensen, L.J. (2014). COMPARTMENTS: unification and visualization of protein subcellular localization evidence. Database 2014, bau012. 10.1093/database/bau012.

45. Aleksander, S.A., Balhoff, J., Carbon, S., Cherry, J.M., Drabkin, H.J., Ebert, D., Feuermann, M., Gaudet, P., Harris, N.L., Hill, D.P., et al. (2023). The Gene Ontology knowledgebase in 2023. Genetics 224, iyad031. 10.1093/genetics/iyad031.

46. Segal, D., Maier, S., Mastromarco, G.J., Qian, W.W., Nabeel-Shah, S., Lee, H., Moore, G., Lacoste, J., Larsen, B., Lin, Z.-Y., et al. (2023). A central chaperone-like role for 14-3-3 proteins in human cells. Mol Cell 83, 974–993.e15. 10.1016/j.molcel.2023.02.018.

47. Chen, T., and Guestrin, C. (2016). XGBoost: A Scalable Tree Boosting System. arXiv. 10.1145/2939672.2939785.

48. Friedman, J.H. (2001). Greedy function approximation: A gradient boosting machine. Ann. Stat. 29. 10.1214/aos/1013203451.

49. Geladaki, A., Britovšek, N.K., Breckels, L.M., Smith, T.S., Vennard, O.L., Mulvey, C.M., Crook, O.M., Gatto, L., and Lilley, K.S. (2019). Combining LOPIT with differential ultracentrifugation for high-resolution spatial proteomics. Nature Communications 10, 331. 10.1038/s41467-018-08191-w.

50. Rath, S., Sharma, R., Gupta, R., Ast, T., Chan, C., Durham, T.J., Goodman, R.P., Grabarek, Z., Haas, M.E., Hung, W.H.W., et al. (2020). MitoCarta3.0: an updated mitochondrial proteome now with sub-organelle localization and pathway annotations. Nucleic Acids Res. 49, D1541– D1547. 10.1093/nar/gkaa1011.

51. King, J.S., Gueho, A., Hagedorn, M., Gopaldass, N., Leuba, F., Soldati, T., and Insall, R.H. (2013). WASH is required for lysosomal recycling and efficient autophagic and phagocytic digestion. Mol. Biol. Cell 24, 2714–2726. 10.1091/mbc.e13-02-0092.

52. McNally, K.E., and Cullen, P.J. (2018). Endosomal Retrieval of Cargo: Retromer Is Not Alone. Trends Cell Biol. 28, 807–822. 10.1016/j.tcb.2018.06.005.

53. Tu, Y., Zhao, L., Billadeau, D.D., and Jia, D. (2020). Endosome-to-TGN Trafficking: Organelle-Vesicle and Organelle-Organelle Interactions. Front. Cell Dev. Biol. 8, 163. 10.3389/fcell.2020.00163.

54. McNally, K.E., Faulkner, R., Steinberg, F., Gallon, M., Ghai, R., Pim, D., Langton, P., Pearson, N., Danson, C.M., Nägele, H., et al. (2017). Retriever is a multiprotein complex for retromer-independent endosomal cargo recycling. Nat. Cell Biol. 19, 1214–1225. 10.1038/ncb3610.

55. Healy, M.D., McNally, K.E., Butkovič, R., Chilton, M., Kato, K., Sacharz, J., McConville, C., Moody, E.R.R., Shaw, S., Planelles-Herrero, V.J., et al. (2023). Structure of the endosomal Commander complex linked to Ritscher-Schinzel syndrome. Cell 186, 2219–2237.e29. 10.1016/j.cell.2023.04.003.

56. Ivanov, P., Kedersha, N., and Anderson, P. (2018). Stress Granules and Processing Bodies in Translational Control. Cold Spring Harb. Perspect. Biol. 11, a032813. 10.1101/cshperspect.a032813.

57. Kirchhausen, T., Owen, D., and Harrison, S.C. (2014). Molecular Structure, Function, and Dynamics of Clathrin-Mediated Membrane Traffic. Cold Spring Harb. Perspect. Biol. 6, a016725. 10.1101/cshperspect.a016725.

58. Mettlen, M., Chen, P.-H., Srinivasan, S., Danuser, G., and Schmid, S.L. (2018). Regulation of Clathrin-Mediated Endocytosis. Annu. Rev. Biochem. 87, 871–896. 10.1146/annurev-biochem-062917-012644.

59. Ghosh, P., Dahms, N.M., and Kornfeld, S. (2003). Mannose 6-phosphate receptors: new twists in the tale. Nat. Rev. Mol. Cell Biol. 4, 202–213. 10.1038/nrm1050.

60. Balderhaar, H.J. kleine, and Ungermann, C. (2013). CORVET and HOPS tethering complexes - coordinators of endosome and lysosome fusion. J Cell Sci 126, 1307–1316. 10.1242/jcs.107805.

61. Khatter, D., Raina, V.B., Dwivedi, D., Sindhwani, A., Bahl, S., and Sharma, M. (2015). The small GTPase Arl8b regulates assembly of the mammalian HOPS complex on lysosomes. J. Cell Sci. 128, 1746–1761. 10.1242/jcs.162651.

62. Guerra, F., and Bucci, C. (2016). Multiple Roles of the Small GTPase Rab7. Cells 5, 34. 10.3390/cells5030034.

63. Kobielak, A., and Fuchs, E. (2004). α-catenin: at the junction of intercellular adhesion and actin dynamics. Nat. Rev. Mol. Cell Biol. 5, 614–625. 10.1038/nrm1433.

64. Stossel, T.P., Condeelis, J., Cooley, L., Hartwig, J.H., Noegel, A., Schleicher, M., and Shapiro, S.S. (2001). Filamins as integrators of cell mechanics and signalling. Nat. Rev. Mol. Cell Biol. 2, 138–145. 10.1038/35052082.

65. Ridley, A.J. (2015). Rho GTPase signalling in cell migration. Curr. Opin. Cell Biol. 36, 103–112. 10.1016/j.ceb.2015.08.005.

66. Consortium, T.U., Bateman, A., Martin, M.-J., Orchard, S., Magrane, M., Ahmad, S., Alpi, E., Bowler-Barnett, E.H., Britto, R., Bye-A-Jee, H., et al. (2022). UniProt: the Universal Protein Knowledgebase in 2023. Nucleic Acids Res. 51, D523–D531. 10.1093/nar/gkac1052.

67. Cooper-Knock, J., Moll, T., Ramesh, T., Castelli, L., Beer, A., Robins, H., Fox, I., Niedermoser, I., Damme, P.V., Moisse, M., et al. (2019). Mutations in the Glycosyltransferase Domain of GLT8D1 Are Associated with Familial Amyotrophic Lateral Sclerosis. Cell Rep. 26, 2298–2306.e5. 10.1016/j.celrep.2019.02.006.

68. Ilina, E.I., Cialini, C., Gerloff, D.L., Garcia-Escudero, M.D., Jeanty, C., Thézénas, M.-L., Lesur, A., Puard, V., Bernardin, F., Moter, A., et al. (2022). Enzymatic activity of glycosyltransferase GLT8D1 promotes human glioblastoma cell migration. iScience 25, 103842. 10.1016/j.isci.2022.103842.

69. Hoek, L.V.D. (2007). Human Coronaviruses: What Do They Cause? Antivir. Ther. 12, 651– 658. 10.1177/135965350701200s01.1.

70. V’kovski, P., Kratzel, A., Steiner, S., Stalder, H., and Thiel, V. (2020). Coronavirus biology and replication: implications for SARS-CoV-2. Nat Rev Microbiol, 1–16. 10.1038/s41579-020-00468-6.

71. Ghosh, S., Dellibovi-Ragheb, T.A., Kerviel, A., Pak, E., Qiu, Q., Fisher, M., Takvorian, P.M., Bleck, C., Hsu, V.W., Fehr, A.R., et al. (2020). β-Coronaviruses Use Lysosomes for Egress Instead of the Biosynthetic Secretory Pathway. Cell 183, 1520–1535.e14. 10.1016/j.cell.2020.10.039.

72. Gordon, D.E., Jang, G.M., Bouhaddou, M., Xu, J., Obernier, K., White, K.M., O’Meara, M.J., Rezelj, V.V., Guo, J.Z., Swaney, D.L., et al. (2020). A SARS-CoV-2 protein interaction map reveals targets for drug repurposing. Nature 583. 10.1038/s41586-020-2286-9.

73. Ogata, H., Goto, S., Sato, K., Fujibuchi, W., Bono, H., and Kanehisa, M. (1999). KEGG: Kyoto Encyclopedia of Genes and Genomes. Nucleic Acids Res. 27, 29–34. 10.1093/nar/27.1.29.

74. Kanehisa, M., Furumichi, M., Sato, Y., Kawashima, M., and Ishiguro-Watanabe, M. (2022). KEGG for taxonomy-based analysis of pathways and genomes. Nucleic Acids Res. 51, D587– D592. 10.1093/nar/gkac963.

75. Wang, R., Simoneau, C.R., Kulsuptrakul, J., Bouhaddou, M., Travisano, K.A., Hayashi, J.M., Carlson-Stevermer, J., Zengel, J.R., Richards, C.M., Fozouni, P., et al. (2021). Genetic Screens Identify Host Factors for SARS-CoV-2 and Common Cold Coronaviruses. Cell 184, 106–119.e14. 10.1016/j.cell.2020.12.004.

76. Martín, C., Fernández-Vega, I., Suárez, J.E., and Quirós, L.M. (2020). Adherence of Lactobacillus salivarius to HeLa Cells Promotes Changes in the Expression of the Genes Involved in Biosynthesis of Their Ligands. Front. Immunol. 10, 3019. 10.3389/fimmu.2019.03019.

77. Prentice, E., Jerome, W.G., Yoshimori, T., Mizushima, N., and Denison, M.R. (2004). Coronavirus Replication Complex Formation Utilizes Components of Cellular Autophagy*. J. Biol. Chem. 279, 10136–10141. 10.1074/jbc.m306124200.

78. Maier, H.J., and Britton, P. (2012). Involvement of Autophagy in Coronavirus Replication. Viruses 4, 3440–3451. 10.3390/v4123440.

79. Min, J.S., Kim, D.E., Jin, Y.-H., and Kwon, S. (2020). Kurarinone Inhibits HCoV-OC43 Infection by Impairing the Virus-Induced Autophagic Flux in MRC-5 Human Lung Cells. J. Clin. Med. 9, 2230. 10.3390/jcm9072230.

80. Lee, S., Yu, Y., Trimpert, J., Benthani, F., Mairhofer, M., Richter-Pechanska, P., Wyler, E., Belenki, D., Kaltenbrunner, S., Pammer, M., et al. (2021). Virus-induced senescence is a driver and therapeutic target in COVID-19. Nature 599, 283–289. 10.1038/s41586-021-03995-1.

81. Kung, Y.-A., Chiang, H.-J., Li, M.-L., Gong, Y.-N., Chiu, H.-P., Hung, C.-T., Huang, P.-N., Huang, S.-Y., Wang, P.-Y., Hsu, T.-A., et al. (2022). Acyl-Coenzyme A Synthetase Long-Chain Family Member 4 Is Involved in Viral Replication Organelle Formation and Facilitates Virus Replication via Ferroptosis. mBio 13, e02717–21. 10.1128/mbio.02717-21.

82. Li, J., Cao, F., Yin, H., Huang, Z., Lin, Z., Mao, N., Sun, B., and Wang, G. (2020). Ferroptosis: past, present and future. Cell Death Dis. 11, 88. 10.1038/s41419-020-2298-2.

83. Jiang, X., Stockwell, B.R., and Conrad, M. (2021). Ferroptosis: mechanisms, biology and role in disease. Nat. Rev. Mol. Cell Biol. 22, 266–282. 10.1038/s41580-020-00324-8.

84. Hou, W., Xie, Y., Song, X., Sun, X., Lotze, M.T., Zeh, H.J., Kang, R., and Tang, D. (2016). Autophagy promotes ferroptosis by degradation of ferritin. Autophagy 12, 1425–1428. 10.1080/15548627.2016.1187366.

85. Gao, M., Monian, P., Pan, Q., Zhang, W., Xiang, J., and Jiang, X. (2016). Ferroptosis is an autophagic cell death process. Cell Res. 26, 1021–1032. 10.1038/cr.2016.95.

86. Chen, X., Yu, C., Kang, R., and Tang, D. (2020). Iron Metabolism in Ferroptosis. Front. Cell Dev. Biol. 8, 590226. 10.3389/fcell.2020.590226.

87. Doll, S., Proneth, B., Tyurina, Y.Y., Panzilius, E., Kobayashi, S., Ingold, I., Irmler, M., Beckers, J., Aichler, M., Walch, A., et al. (2017). ACSL4 dictates ferroptosis sensitivity by shaping cellular lipid composition. Nat. Chem. Biol. 13, 91–98. 10.1038/nchembio.2239.

88. Mancias, J.D., Wang, X., Gygi, S.P., Harper, J.W., and Kimmelman, A.C. (2014). Quantitative proteomics identifies NCOA4 as the cargo receptor mediating ferritinophagy. Nature 509, 105–109. 10.1038/nature13148.

89. Hung, V., Lam, S.S., Udeshi, N.D., Svinkina, T., Guzman, G., Mootha, V.K., Carr, S.A., and Ting, A.Y. (2017). Proteomic mapping of cytosol-facing outer mitochondrial and ER membranes in living human cells by proximity biotinylation. eLife 6. 10.7554/elife.24463.

90. Brunner, A., Thielert, M., Vasilopoulou, C., Ammar, C., Coscia, F., Mund, A., Hoerning, O.B., Bache, N., Apalategui, A., Lubeck, M., et al. (2022). Ultra-high sensitivity mass spectrometry quantifies single-cell proteome changes upon perturbation. Mol. Syst. Biol. 18, e10798. 10.15252/msb.202110798.

91. Crook, O.M., Davies, C.T.R., Breckels, L.M., Christopher, J.A., Gatto, L., Kirk, P.D.W., and Lilley, K.S. (2022). Inferring differential subcellular localisation in comparative spatial proteomics using BANDLE. Nat. Commun. 13, 5948. 10.1038/s41467-022-33570-9.

92. Hashimoto, Y., Sheng, X., Murray-Nerger, L.A., and Cristea, I.M. (2020). Temporal dynamics of protein complex formation and dissociation during human cytomegalovirus infection. Nat Commun 11, 806. 10.1038/s41467-020-14586-5.

93. Marx, V. (2021). Method of the Year: spatially resolved transcriptomics. Nat. Methods 18, 9–14. 10.1038/s41592-020-01033-y.

94. McGinnis, C.S., Patterson, D.M., Winkler, J., Conrad, D.N., Hein, M.Y., Srivastava, V., Hu, J.L., Murrow, L.M., Weissman, J.S., Werb, Z., et al. (2019). MULTI-seq: sample multiplexing for single-cell RNA sequencing using lipid-tagged indices. Nat. Methods 16, 619–626. 10.1038/s41592-019-0433-8.

95. Sunshine, S., Puschnik, A.S., Replogle, J.M., Laurie, M.T., Liu, J., Zha, B.S., Nuñez, J.K., Byrum, J.R., McMorrow, A.H., Frieman, M.B., et al. (2023). Systematic functional interrogation of SARS-CoV-2 host factors using Perturb-seq. Nat. Commun. 14, 6245. 10.1038/s41467-023-41788-4.

96. Cox, J., and Mann, M. (2008). MaxQuant enables high peptide identification rates, individualized p.p.b.-range mass accuracies and proteome-wide protein quantification. Nat Biotechnol 26, 1367–1372. 10.1038/nbt.1511.

97. Cox, J., Hein, M.Y., Luber, C.A., Paron, I., Nagaraj, N., and Mann, M. (2014). Accurate proteome-wide label-free quantification by delayed normalization and maximal peptide ratio extraction, termed MaxLFQ. Mol Cell Proteom Mcp 13, 2513–2526. 10.1074/mcp.m113.031591.

98. Wolf, F.A., Angerer, P., and Theis, F.J. (2018). SCANPY: large-scale single-cell gene expression data analysis. Genome Biol 19, 15. 10.1186/s13059-017-1382-0.

99. Kuleshov, M.V., Jones, M.R., Rouillard, A.D., Fernandez, N.F., Duan, Q., Wang, Z., Koplev, S., Jenkins, S.L., Jagodnik, K.M., Lachmann, A., et al. (2016). Enrichr: a comprehensive gene set enrichment analysis web server 2016 update. Nucleic Acids Res. 44, W90–W97. 10.1093/nar/gkw377.

100. Mitchell, R., and Frank, E. (2017). Accelerating the XGBoost algorithm using GPU computing. PeerJ Comput. Sci. 3, e127. 10.7717/peerj-cs.127.

101. Vairetti, C., Assadi, J.L., and Maldonado, S. (2023). Efficient Hybrid Oversampling and Intelligent Undersampling for Imbalanced Big Data Classification. arXiv. 10.48550/arxiv.2310.05789.

102. 102. Snoek, J., Larochelle, H., and Adams, R.P. (2012). Practical Bayesian Optimization of Machine Learning Algorithms. arXiv. 10.48550/arxiv.1206.2944.

